# The impact of physical activity on physical performance, mitochondrial bioenergetics, ROS production and calcium handling across the human adult lifespan

**DOI:** 10.1101/2024.07.09.602758

**Authors:** Marina Cefis, Vincent Marcangeli, Rami Hammad, Jordan Granet, Jean-Philippe Leduc-Gaudet, Pierrette Gaudreau, Caroline Trumpff, Hannah Huang, Martin Picard, Mylène Aubertin-Leheudre, Marc Bélanger, Richard Robitaille, José A. Morais, Gilles Gouspillou

## Abstract

Aging-related muscle atrophy and weakness contribute to loss of mobility, falls and disability. Mitochondrial dysfunction is widely considered a key contributing mechanism to muscle aging. However, mounting evidence position physical activity as a confounding factor, making unclear whether muscle mitochondria accumulate bona fide defects with aging. To disentangle aging from physical activity-related mitochondrial adaptations, we functionally profiled skeletal muscle mitochondria in 51 inactive and 88 active men aged 20-93. Physical activity status conferred partial protection against age-related decline in physical performance. A trend for reduced muscle mitochondrial respiration with aging was observed in inactive but not in active participants, indicating that aging *per se* does not alter mitochondrial respiratory capacity. Mitochondrial reactive oxygen species (ROS) production was unaffected by aging and active participants displayed higher ROS production. In contrast, mitochondrial calcium retention capacity decreased with aging regardless of physical activity status and correlated with muscle mass, performance and the stress-responsive metabokine GDF15. Targeting mitochondrial calcium handling may hold promise for treating aging-related muscle impairments.

## Introduction

One of the most significant changes associated with normal aging is a progressive loss of muscle mass and function ^1–3^. These age-related skeletal muscle alterations greatly contribute to reduced functional capacities, mobility impairment, falls, and physical frailty in older adults, dramatically impairing their quality of life ^4,5^. Underscoring the magnitude of the impact of aging-related muscle dysfunction, it is estimated that the prevalence of sarcopenia (i.e., the aging-related loss of muscle mass and function) increases from 14% in 65 to 70 year old, to over 50% in those above 80 years of age ^6^, underscoring the need for studies in individuals across the whole adult lifespan. The number of people around the world aged over 60 years was estimated at 1 billion in the year 2020. This figure is expected to more than double by 2025 (https://www.who.int/health-topics/ageing). Even with a conservative estimate of its prevalence, sarcopenia currently affects over 50 million people and will reach over 200 millions in the next 30 years ^6^. Furthermore, the direct health care costs attributable to sarcopenia and its sequela are considerable, estimated at 18.5 billion dollars for 2000 alone in the US ^7^, a figure certainly much higher today. Hence, unraveling the mechanisms underlying the aging-related loss of muscle mass and function to promote the development of effective therapeutic interventions is a major challenge in health research.

Amongst the mechanisms proposed to explain the aging-related alteration in skeletal muscle biology, the accumulation of mitochondrial dysfunction gained a lot of traction in the past few decades. Impaired mitochondrial function is often depicted as a hallmark of muscle aging with a reduction in mitochondrial respiration and an increase in mitochondrial reactive oxygen species (ROS) production commonly portrayed as key contributing mechanisms ^8–16^. Although debated ^17^, the aging-related accumulation of mitochondrial DNA mutations has been proposed as an underlying mechanism to reduced mitochondrial respiratory function with aging ^18–21^. Impaired mitochondrial calcium handling and altered function of the mitochondrial permeability transition pore (mPTP), and potential downstream activation of apoptotic and proteolytic pathways, have also been proposed as potential contributors to skeletal muscle aging ^8,17,22^. However, age-related alterations in skeletal muscle mitochondrial biology are not universal findings, and mounting evidence indicate that habitual physical activity is a likely confounding factor when interpreting the impact of aging ^14,15,22–27^. Most studies conducted in humans also focus on comparisons between young *vs* older adults, potentially missing changes that could occur during middle age. Those who did include middle-aged adults mainly focused on mitochondrial bioenergetics (i.e. respiration or ATP production) and did not investigate the impact of habitual physical activity (i.e. by comparing inactive *vs* active individuals) ^28,29^.

To address these gaps in knowledge, we conducted a cross-sectional study with deep muscle mitochondrial phenotyping in 51 inactive and 88 active men with ages ranging from 20 to 93 years, divided in age groups to tease out when significant changes in parameters may occur. All participants were deeply phenotyped using a comprehensive battery of physical tests. Body and skeletal muscle composition were assessed using dual-energy X-ray absorptiometry (DXA) and peripheral quantitative computed tomography (pQCT), respectively. Vastus lateralis muscle biopsies were performed to assess various aspects of mitochondrial function, including mitochondrial respiration, ROS production (assessed using H2O2 emission as a surrogate), and calcium handling in permeabilized myofibers. Group and correlation analyses were performed to assess the impact of aging and physical activity status on physical performance, mitochondrial bioenergetics, ROS production and calcium handling and to investigate potential associations between various aspects of mitochondrial function and measures of physical performance, muscle mass and muscle power.

## Results

### Baseline participant characteristics

One hundred and thirty-nine men aged between 20 to 93 years old, all living independently in the community and in overall good physical health, were enrolled in the present study. To assess the impact of aging and habitual physical activity, analyses were conducted either by grouping participants according to their age and physical activity status or were performed using age as a continuous variable. For group analyses, participants were divided in the following age groups: young middle age (20-39 yo), mature middle age (40-59yo), young older adults (60-69 yo) and older adults (70+ yo) and sub-divided in inactive and active subgroups according to their physical activity status (i.e physically inactive vs active).

Anthropometrics characteristics and body composition data as well as time practicing physical activity are summarized in Table 1. No effect of aging on body mass was observed (Table 1). An effect of physical activity status on body mass, mainly driven by participants in the 40-59 and 70+ groups, was observed and indicates that overall, physically active participants had lower body mass compared to their inactive counterparts (Table 1). Body mass index (BMI) significantly increased with aging, an effect attenuated by physical activity status (Table 1). Total fat mass also increased with aging. A significant effect of physical activity status was observed for fat mass indicative of lower fat mass in physically active participants across all age groups (Table 1). Lean mass relative to body mass was significantly reduced with aging. A significant effect of physical activity status was observed for relative lean mass, indicative of higher relative lean mass in physically active participants across age groups (Table 1). In line with data on relative lean mass. Absolute lean mass was significantly reduced with aging, although only a trend for an effect of physical activity was observed (Table 1). Aging was associated with a reduction in the number of daily steps (Table 1). By design, daily steps, metabolic equivalent of tasks (METs) and self-reported amount of physical activity were higher in the active participants (Table 1). Details on the type of physical activity (time spent performing aerobic and/or resistance training) performed by all participants are provided in Table 1. No significant impact of aging on overall physical activity, as well as time spent performing aerobic and/or resistance training was observed in our cohort (Table 1). However, strong trends for decrease in total physical activity as well as the time spent performing resistance training were noticed in our physically active participants (Table 1).

**Table 1:** Participant characteristics.

|  | GROUP | INACTIVE |  |  |  | ACTIVE |  |  |  | 2-WAY ANOVA (p values) |  |  |
| --- | --- | --- | --- | --- | --- | --- | --- | --- | --- | --- | --- | --- |
|  |  | 20-39<br>(n=15) | 40-59<br>(n=14) | 60-69<br>(n=9) | 70+<br>(n=13) | 20-39<br>(n=29) | 40-59<br>(n=25) | 60-69<br>(n=18) | 70+<br>(n=16) | Age | PA | Interaction |
| Characteristics & body composition | Age (years) | 30.3 ± 4.1 | 52.1 ± 4.9 | 64.1 ± 1.5 | 79.1 ± 7.8 | 29.7 ± 5.5 | 49.6 ± 6.9 | 63.7 ± 2.7 | 75.9 ± 4.6 | <0.001 | 0.087 | 0.676 |
|  | Weight (kg) | 78.3 ± 13.2 | 87.9 ± 14.8 | 81.1 ± 12.5 | 81.5 ± 8.9 | 78.8 ± 12.0 | 75.7 ± 10.3 | 81.0 ± 10.8 | 76.4 ± 12.7 | 0.613 | 0.049 | 0.105 |
|  | BMI (kg.m<sup>-2</sup>) | 24.2 ± 3.6 | 28.3 ± 3.5 | 27.4 ± 3.8 | 28.6 ± 2.5 | 25.3 ± 3.7 | 25.3 ± 3.3 | 26.7 ± 3.5 | 25.6 ± 3.3 | 0.013 | 0.026 | 0.033 |
|  | Fat mass (%) | 27.11 ± 7.8 | 31.7 ± 5.6 | 29.3 ± 7.2 | 32.3 ± 5.2 | 17.8 ± 7.9 | 21.4 ± 7.3 | 27.2 ± 7.9 | 27.2 ± 8.0 | <0.001 | <0.001 | 0.117 |
|  | Lean mass (%) | 69.9 ± 7.2 | 65.6 ± 5.3 | 68.1 ± 6.7 | 65.0 ± 4.9 | 78.6 ± 7.4 | 75.3 ± 7.0 | 69.9 ± 7.6 | 69.7 ± 7.6 | <0.001 | <0.001 | 0.111 |
|  | Lean mass (kg) | 54.2 ± 6.9 | 57.3 ± 8.6 | 54.6 ± 4.9 | 52.7 ± 4.4 | 61.4 ± 7.3 | 56.6 ± 5.4 | 56.0 ± 4.7 | 52.6 ± 5.8 | 0.007 | 0.088 | 0.026 |
| Physical activity level | Steps per day (n) | 5830 ± 2107 | 6129 ± 2043 | 6781 ± 1811 | 3976 ± 2276 | 10995 ± 4000 | 11743 ± 5346 | 8427 ± 4394 | 5250 ± 2701 | <0.001 | <0.001 | 0.081 |
|  | METs (kcal.kg<sup>-1</sup>.h<sup>-1</sup>) | 1.39 ± 0.14 | 1.26 ± 0.14 | 1.22 ± 0.20 | 1.07 ± 0.14 | 1.67 ± 0.36 | 1.62 ± 0.43 | 1.31 ± 0.30 | 1.13 ± 0.22 | <0.001 | 0.001 | 0.211 |
|  | Physical activity total (min.week<sup>-1</sup>) | 38.3 ± 45.8 | 37.0 ± 42.9 | 27.5 ± 55.4 | 41.8 ± 46.4 | 635.9 ± 381.9 | 355.7 ± 288.5 | 371.1 ± 247.2 | 377.2 ± 269.8 | 0.051 | <0.001 | 0.057 |
|  | Aerobic training (min.week<sup>-1</sup>) | 38.3 ± 45.8 | 32.7 ± 43.4 | 27.5 ± 55.4 | 19.6 ± 38.9 | 413.7 ± 383.9 | 257.7 ± 260.9 | 301.1 ± 249.8 | 220.8 ± 156.8 | 0.275 | <0.001 | 0.410 |
|  | Resistance training (min week<sup>-1</sup>) | 0 ± 0 | 4.3 ± 16.0 | 0 ± 0 | 22.2 ± 35.3 | 222.2 ± 227.2 | 97.8 ± 126.8 | 70.0 ± 78.2 | 156.4 ± 159.8 | 0.094 | <0.001 | 0.096 |
| Blood biochemistry | Glucose (mmol.l<sup>-1</sup>) | 5.1 ± 0.6 | 5.2 ± 0.4 | 5.8 ± 0.9 | 5.6 ± 0.7 | 4.8 ± 0.3 | 5.0 ± 0.4 | 5.4 ± 0.8 | 5.5 ± 1.0 | <0.001 | 0.037 | 0.928 |
|  | Insulin (pmol.l<sup>-1</sup>) | 61.0 ± 60.4 | 64.2 ± 37.5 | 61.6 ± 25.6 | 52.1 ± 39.4 | 32.2 ± 20.1 | 39.0 ± 33.6 | 48.1 ± 37.1 | 44.1 ± 22.8 | 0.817 | 0.004 | 0.614 |
|  | QUICKI | 0.35 ± 0.04 | 0.35 ± 0.03 | 0.34 ± 0.03 | 0.35 ± 0.03 | 0.39 ± 0.05 | 0.38 ± 0.03 | 0.38 ± 0.08 | 0.36 ± 0.03 | 0.309 | <0.001 | 0.588 |
|  | HOMA-IR | 2.4 ± 2.6 | 2.5 ± 1.6 | 2.8 ± 1.5 | 2.2 ± 1.7 | 1.2 ± 0.8 | 1.5 ± 1.3 | 2.1 ± 2.1 | 1.8 ± 1.1 | 0.479 | 0.005 | 0.703 |
|  | Fructosamine (μmol.l<sup>-1</sup>) | 233.9 ± 26.4 | 223.6 ± 24.9 | 235.7 ± 26.7 | 254.3 ± 35.3 | 232.1 ± 21.6 | 229.4 ± 22.8 | 235.4 ± 34.3 | 238.5 ± 35.9 | 0.052 | 0.565 | 0.506 |
|  | Fructosamine per protein (μmol l<sup>-1</sup>.g<sup>-1</sup>) | 3.2 ± 0.4 | 3.2 ± 0.4 | 3.2 ± 0.4 | 3.7 ± 0.5 | 3.3 ± 0.3 | 3.3 ± 0.3 | 3.4 ± 0.5 | 3.5 ± 0.4 | 0.009 | 0.608 | 0.434 |
|  | Total cholesterol (mmol.l<sup>-1</sup>) | 4.4 ± 0.8 | 5.1 ± 1.1 | 4.9 ± 0.8 | 4.0 ± 0.9 | 4.3 ± 0.8 | 4.8 ± 0.8 | 4.6 ± 1.0 | 4.9 ± 0.9 | 0.057 | 0.825 | 0.032 |
|  | HDL cholesterol (mmol.l<sup>-1</sup>) | 1.3 ± 0.2 | 1.3 ± 0.3 | 1.4 ± 0.2 | 1.4 ± 0.4 | 1.4 ± 0.3 | 1.5 ± 0.3 | 1.4 ± 0.3 | 1.4 ± 0.4 | 0.654 | 0.188 | 0.366 |
|  | LDL cholesterol (mmol.l<sup>-1</sup>) | 2.7 ± 0.7 | 2.7 ± 0.8 | 2.9 ± 0.8 | 2.0 ± 0.7 | 2.5 ± 0.8 | 2.7 ± 0.7 | 2.6 ± 0.8 | 2.8 ± 0.9 | 0.343 | 0.544 | 0.052 |
|  | Triglycerides (mmol.l<sup>-1</sup>) | 1.0 ± 0.5 | 2.2 ± 1.4 | 1.3 ± 0.3 | 1.3 ± 0.5 | 1.0 ± 0.4 | 1.1 ± 0.5 | 1.1 ± 0.5 | 1.5 ± 0.7 | 0.001 | 0.038 | <0.001 |
|  | C-reactive protein (mg.l<sup>-1</sup>) | 1.3 ± 1.4 | 1.8 ± 1.1 | 2.6 ± 3.2 | 1.2 ± 0.7 | 0.6 ± 0.4 | 1.5 ± 2.6 | 1.4 ± 0.9 | 2.4 ± 3.5 | 0.164 | 0.480 | 0.110 |
|  | Proteins (g.l<sup>-1</sup>) | 72.9 ± 2.8 | 69.4 ± 2.9 | 72.6 ± 2.4 | 69.9 ± 2.9 | 71.0 ± 2.9 | 69.2 ± 3.9 | 68.9 ± 2.9 | 67.5 ± 4.5 | 0.001 | 0.003 | 0.337 |
|  | GDF-15 (pg.ml<sup>-1</sup>) | 361 ± 116 | 571 ± 136 | 631 ± 305 | 1857 ± 990 | 329 ± 93 | 475 ± 99 | 780 ± 404 | 1393 ± 714 | <0.001 | 0.152 | 0.063 |

BMI, Body Mass Index; METs, Metabolic Equivalent of Tasks, QUICKI, Quantitative Insulin Sensitivity Check Index, HOMA-IR, Homeostatic Model Assessment for Insulin Resistance; HDL, High-Density Lipoprotein; LDL, Low-Density Lipoprotein; GDF15, Growth Differentiation Factor 15. Data are expressed as mean ± SD. p<0.05 was considered statistically significant.

To further characterize our cohort of participants, various blood biochemical analyses were performed. As shown in Table 1, significant effects of aging and physical activity status were observed for circulating blood glucose concentration assessed in fasted state, indicative of an increase of fasted glycemia with aging and a lower fasted glycemia in physically active individuals (Table 1). No effect of aging on circulating insulin levels was observed. However, physically active individuals displayed lower circulating insulin levels (Table 1). While no effect of aging was found for the Quantitative Insulin Sensitivity Check Index (QUICKI) and the Homeostatic Model Assessment of Insulin Resistance (HOMA-IR), 2 indexes of whole-body insulin sensitivity, physically active individuals displayed greater QUICK-I values and lower HOMA-I values, both indicative of greater whole body insulin sensitivity in active participants (Table 1). Circulating fructosamine levels significantly increase with aging, without protection from physical activity status (Table 1). A trend for an increase in total cholesterol with aging was observed, without protection from physical activity status (Table 1). No impact of aging or physical activity status on HDL and LDL cholesterol was observed. Significant effects of aging and physical activity status on circulating triglycerides levels were overserved, indicative of an increase in circulating triglycerides levels with aging, with a protection conferred by the physical activity status (Table 1). However, the effect of physical activity seemed mainly driven by the inactive and active participants in the 40-59 groups (Table 1). No effect of aging or physical activity on the circulating levels of C-reactive protein was observed (Table 1). Significant effects of aging and physical activity on the total protein content in serum were observed, indicative of a decrease in total protein content in serum with aging and lower total protein content in active vs inactive participants (Table 1). Taken altogether, these results indicate that aging is associated with alterations in glucose handling, insulin sensitivity and triglycerides levels, changes that appear partly prevented by physical activity status. However, it should be noted that average values displayed in Table 1 for most circulating markers fall well within normal ranges, indicating that both cohorts of inactive and active participants were overall in good general health.

### The impact of aging and physical activity status on physical function

All participants underwent a comprehensive battery of tests to assess their physical function. As shown in Fig. 1A-H, performance on the 6-minute walk test (Fig.1A-B), step test (Fig.1C-D), sit to stand tests (Fig.1E-F and Fig. S1A-B) and fast-paced timed up and go (Fig.1G-H) were significantly decreased with aging. Significant effects of physical activity status were found for all tested functional capacities, indicating that physically active participants displayed greater functional capacities compared to their inactive counterparts (Fig.1A, C, E, G). Taken altogether, our data show that aging negatively impacts physical function and highlight that physical activity increases physical function in young to older men.

**Figure 1:**
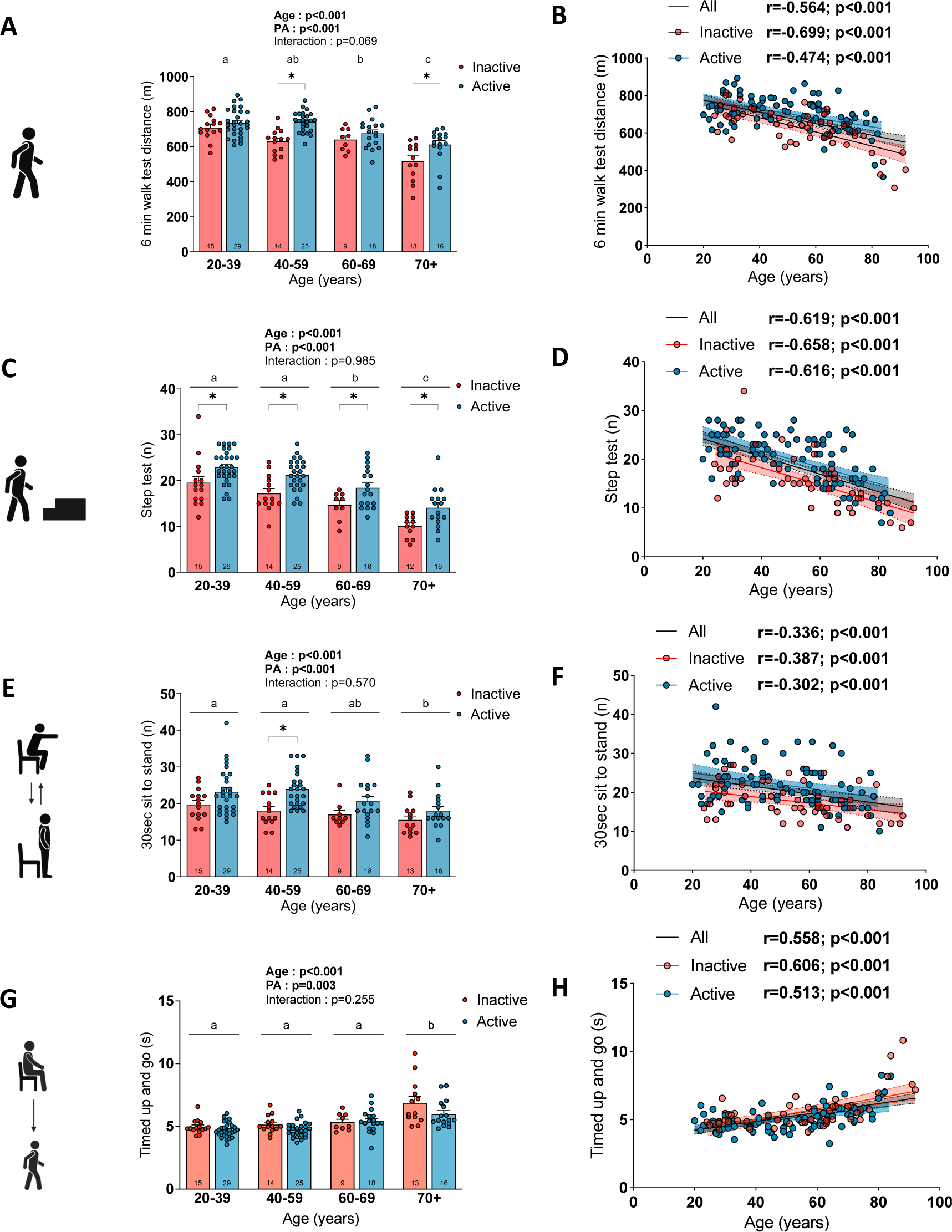
The impact of aging and physical activity status on physical function. Quantification of the performance at the (A, B) 6 min walk test, (C, D) step test, (E,F) sit to stand test (number of repetitions performed in 30s) and (F, G) timed up and go test in inactive and active participants. For group analyses, results of the two-way ANOVA are displayed above each bar graph. Tukey post hoc tests were performed to test differences between age groups. Differences between inactive and active participants within each age group was assessed using multiple bilateral t-tests with FDR correction. Groups that do not share the same letter are significantly different. Linear regressions were performed to assess associations between age and variables of interest in the entire cohort (all) and for inactive and active participants separately. Pearson correlation coefficient (r) and p-values are displayed above each scatter plot. For two-way ANOVA, post hoc testing, and regression analyses, p<0.05 was considered statistically significant. For FDR analyses, q<0.05 was considered statistically significant. *=q<0.05. Icons in panels A, C, E and G were created with BioRender.com.

### The impact of aging and physical activity status on muscle strength and power

We next evaluated the impact of aging and physical activity status on maximal isometric knee extension strength and maximal lower limb power. As shown in Figure 2A-D and Fig. S1C-J, both absolute and relative (to thigh lean mass and body mass) isometric knee extension strength and lower limb extension power were significantly reduced with aging. No significant effect of physical activity status on absolute isometric knee extension strength and absolute lower limb extension power were observed (Fig S1C,D,G,H). Similarly, no significant effect of physical activity status on isometric knee extension strength and lower limb extension power normalized to thigh lean mass were observed (Fig S1E,F,I,J). However, there were significant physical activity status effects on isometric knee extension strength and relative lower limb extension power normalized to body mass (Fig. 2A-D), indicating greater muscle strength and power relative to body mass in physically active compared to physically inactive individuals. Taken altogether, these data extend the literature showing that muscle strength and power decline with aging and highlight the beneficial effect of physical activity status on muscle strength and power relative to body mass.

**Figure 2.**
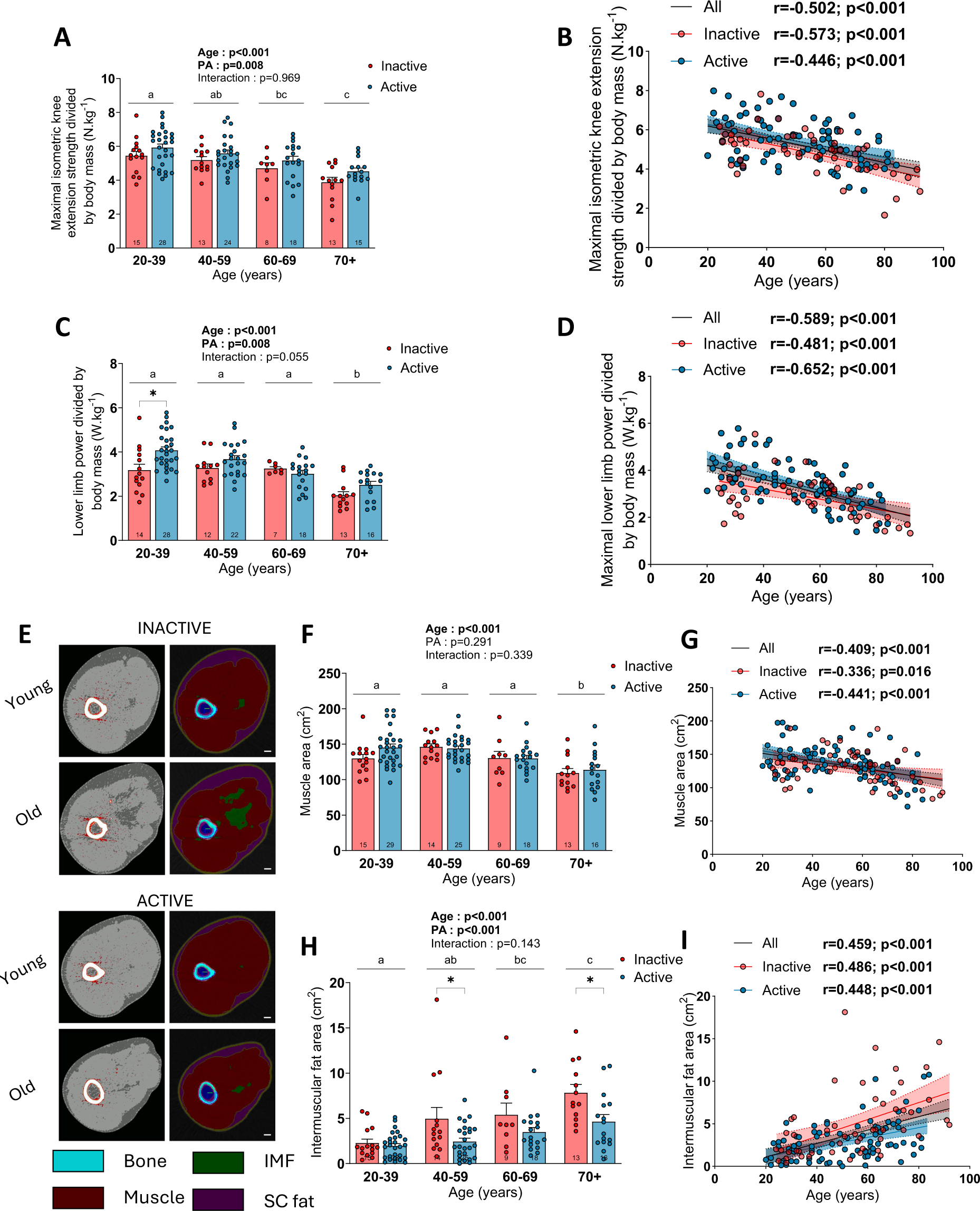
The impact of aging and physical activity status on muscle strength, power and composition. (A, B) Quantification of the maximal isometric knee extension strength relative to body mass in inactive and active participants. (C,D) Quantification of the maximal lower limb power relative to body mass in inactive and active participants. (E) Representative pQCT images of the thigh of young and old inactive and active participants. Quantification of the (F-G) muscle cross-sectional area and (H-I) intermuscular fat area in inactive and active participants. For group analyses, results of the two-way ANOVA are displayed above each bar graph. Tukey post hoc tests were performed to test differences between age groups. Differences between inactive and inactive participants within each age group was assessed using multiple bilateral t-tests with FDR correction. Groups that do not share the same letter are significantly different. Linear regressions were performed to assess associations between age and variables of interest in the entire cohort (all) and for inactive and active participants separately. Pearson correlation coefficient (r) and p-values are displayed above each scatter plot. For two-way ANOVA, post hoc testing, and regression analyses, p<0.05 was considered statistically significant. For FDR analyses, q<0.05 was considered statistically significant. *=q<0.05.

### The impact of aging and physical activity status on skeletal muscle composition

To gain insight into the impact of aging and physical activity on skeletal muscle mass and composition, images of the thigh were collected using pQCT scan. Images were acquired at the lower third of the thigh (Fig.2E) and used to quantify muscle area, subcutaneous and intermuscular fat. Segmental analyses of DXA scan images were also performed to assess thigh lean mass. As shown in Figure 2F-I, muscle area significantly decreased with aging while intermuscular fat area increased. In line with these findings, thigh lean mass assessed using DXA also decreased with aging (Fig. S2A-D). There was no age effect on subcutaneous fat area in our global population (Fig. S2E), although a significant negative correlation was found between age and subcutaneous fat area in inactive participants (Fig. S2F). Interestingly, there was no effect of physical activity status on muscle area and thigh lean mass (Fig. 2F and Fig. S2A). However, there was an effect of physical activity status for intermuscular and subcutaneous fat areas (Fig. 2H and S2E), indicative of lower intermuscular and subcutaneous fat in physically active individuals. While young active and inactive participants displayed comparable intermuscular fat, older active groups displayed lower intermuscular fat (Fig. 2H), suggesting that physical activity exert a protective effect against the aging-related increase in intermuscular fat. In contrast, the lower subcutaneous fat content in physically active individuals was mainly explained by lower subcutaneous fat in young active vs inactive individuals, while older active and inactive individuals displayed comparable subcutaneous fat content (Fig. S2E). There was also an effect of physical activity status when thigh lean mass data were expressed relative to body mass, indicating that physically active individuals displayed greater thigh lean mass relative to their overall body mass (Fig. S2C-D).

A triple myosin heavy chain (MHC) immunolabelling was next performed to assess muscle fiber size and type on vastus lateralis muscle cross-sections (Fig. 3A and S3A). As shown in Figure 3B, neither age nor physical activity status affected the overall muscle fiber cross-sectional area (CSA). No significant impact of aging was observed for the proportion of type I (Fig.3C-D). However, a significant effect of physical activity status was found for the proportion of type I fibers (Fig. 3C), indicative higher proportion of type I fibers in active individuals. Somewhat in line with data published by Lexell et al.^30^, a trend for an increase in the proportion of type I fibers with age was observed in inactive participants (Fig.3D). However, no such trend was observed in active individuals. Taken altogether, these data on type I fibers suggest that active young individuals display a greater proportion of type I fiber that is not affected by aging. In contrast, inactive young individuals display a lower proportion of type I fibers, which appears to progressively increase with aging. A significant effect of age was observed for the proportion of type IIa fibers, indicative of a progressive decline in the proportion of type IIa fibers with aging (Fig.3E-F). No effect of physical activity status on the proportion of type IIa fibers was observed. No pure type IIx fibers were found in the present study, consistent with a recent report indicating that pure type IIx fibers are extremely rare in human skeletal muscles ^31,32^. All fibers positive for type IIx MHC also co-expressed type IIa MHC. A trend for an aging effect and a significant physical activity status effect were observed for the proportion of type IIa/IIx fibers in our group analyses (Fig.3 G). When analyzed with age as a continuous variable, a significant positive correlation between age and the proportion of type IIa/IIx fibers was observed when all participants were pooled together and when active participants were analyzed separately (Fig. 3H). However, no correlation between age and the proportion of type IIa/IIx was found in inactive participants (Fig. 3H). Taken altogether, these data suggest that while young active individuals display a lower proportion of type IIa/IIx fibers vs inactive individuals, the proportion of these fibers only increases in active participants. No effect of aging nor physical activity status was observed for the proportion of the hybrid type I/IIa fibers (Fig.S3 B-C).

**Figure 3.**
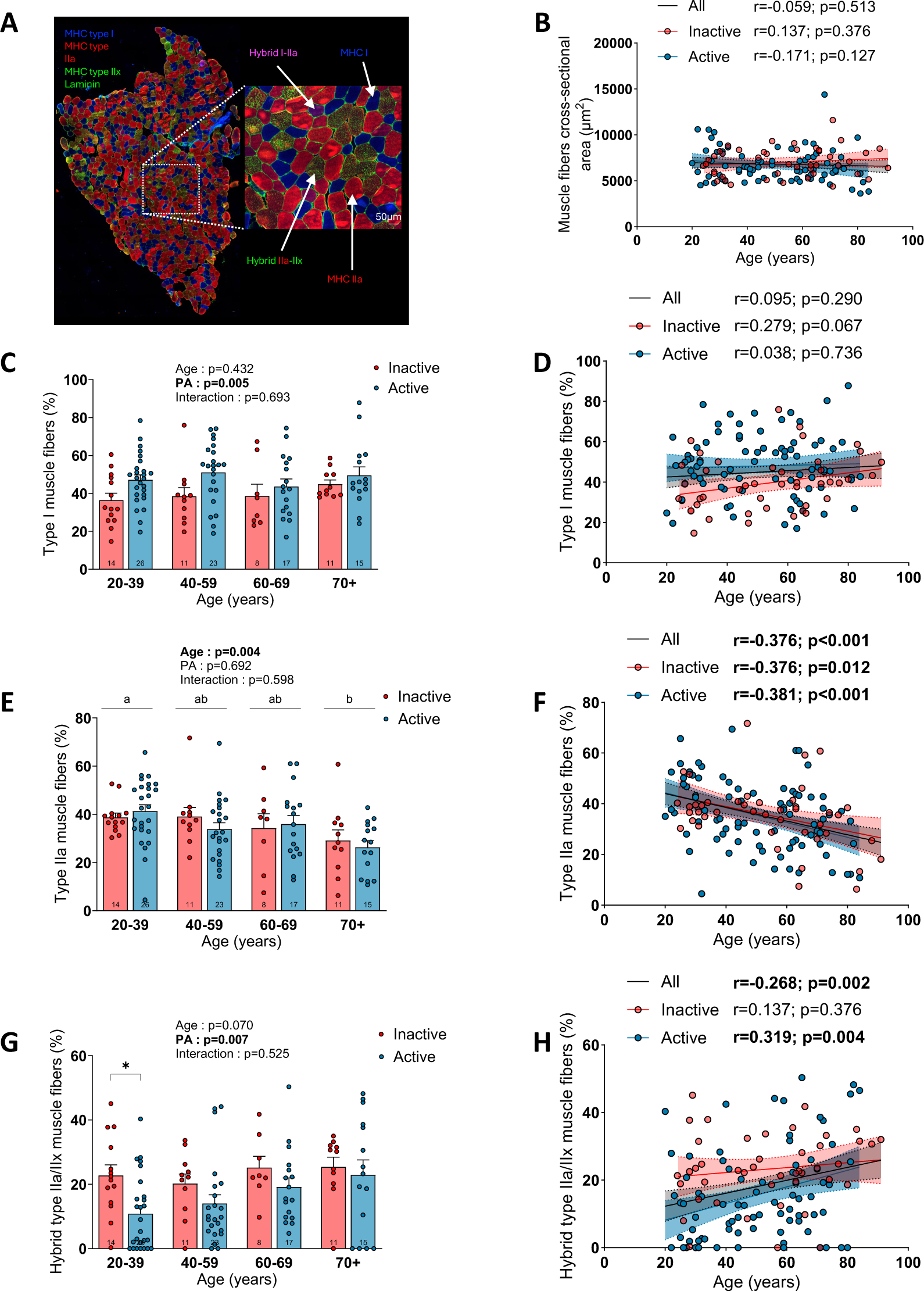
The impacts of aging and physical activity status on muscle fiber type, size and proportion. (A) Representative triple MHCs (MHC type I: blue; MHC type IIa: red; MHC type IIx: green) and laminin (green) immunolabeling performed on muscle cross sections. (B) Quantifications of the overall myofiber cross-sectional area in inactive and active participants. Quantification of the proportion of type I (C,D), IIa (E,F) and IIa/IIx (G,H) myofibers in inactive and active participants. For group analyses, results of the two-way ANOVA are displayed above each bar graph. Tukey post hoc tests were performed to test differences between age groups. Differences between inactive and active participants within each age group was assessed using multiple bilateral t-tests with FDR correction. Groups that do not share the same letter are significantly different. Linear regressions were performed to assess associations between age and variables of interest in the entire cohort (all) and for inactive and active participants separately. Pearson correlation coefficient (r) and p-values are displayed above each scatter plot. For two-way ANOVA, post hoc testing, and regression analyses, p<0.05 was considered statistically significant. For FDR analyses, q<0.05 was considered statistically significant. *=q<0.05.

### The impact of aging and physical activity status on skeletal muscle mitochondrial bioenergetics

As shown in Figure 4A, we prepared permeabilized myofibers from muscle biopsies to perform *ex vivo* high resolution fluorespirometry analysis to assess mitochondrial respiration. State II respiration rate was evaluated using complex I substrates (Glutamate(G)+Malate(M)) of the mitochondrial electron transport chain. State II (ADP-restricted) respiration rate supported by complex I substrates was reduced with aging (Fig. S4A). Physically active participants displayed higher state II respiration rates compared to their inactive counterparts. In contrast, state III (maximal ADP-stimulated) respiration rates supported by complex I (G+M) and complex I + II substrates (G+M+Succinate) were not altered with aging and were significantly higher in active *vs* inactive participants (Fig.4B-C and Fig. S4B). Interestingly, when data were analyzed with age as a continuous variable, a trend for a decrease in maximal respiration was observed for the overall population and for inactive, but not for active participants (Fig. 4C). In inactive participants, a population for which accelerometers provides reliable estimates of physical activity (accelerometer such as the one used in this study largely underestimates the physical activity of participants engaged in physical activities such as swimming, biking, hockey, or resistance training), a significant positive correlation between the number of daily steps and mitochondrial respiration was observed (Fig. S4C). The acceptor control ratio (ACR), an index of the oxidative phosphorylation coupling efficiency (calculated by dividing state III by state II respiration rates) was significantly increased with aging (Fig. S4D). No impact of physical activity status on the ACR was observed (Fig. S4D).

**Figure 4.**
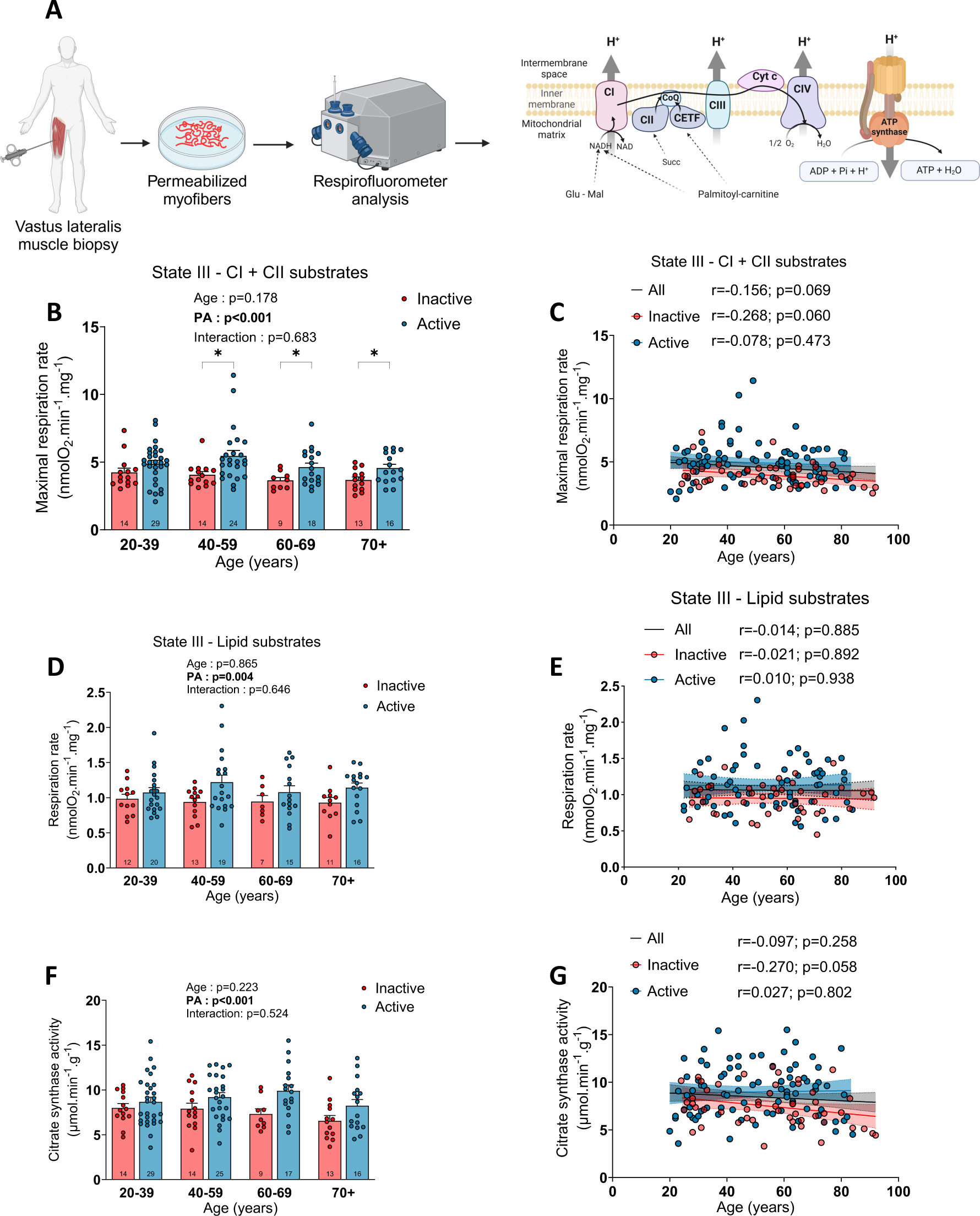
The impact of aging and physical activity status on skeletal muscle mitochondrial respiration and citrate synthase activity. A) Schematic representation of the experimental design used to assess mitochondrial respiration in permeabilized myofibers. (B,C) State III (ADP stimulated) respiration rate driven by complex I and II substrates (Glutamate + Malate + Succinate) in inactive and active participants. (D,E) State III (ADP stimulated) respiration rates driven by lipid substrates (palmitoyl-L-carnitine + malate) in inactive and active participants. (F,G) Citrate synthase activity in inactive and active participants. For group analyses, results of the two-way ANOVA are displayed above each bar graph. Tukey post hoc tests were performed to test differences between age groups. Differences between inactive and active participants within each age group was assessed using multiple bilateral t-tests with FDR correction. Groups that do not share the same letter are significantly different. Linear regressions were performed to assess associations between age and variables of interest in the entire cohort (all) and for inactive and active participants separately. Pearson correlation coefficient (r) and p-values are displayed above each scatter plot. For two-way ANOVA, post hoc testing, and regression analyses, p<0.05 was considered statistically significant. For FDR analyses, q<0.05 was considered statistically significant. *=q<0.05. Panel A was created with BioRender.com.

We also assessed mitochondrial fatty acid oxidation capacity by recording state II and III respiration supported by palmitoyl-L-carnitine (P) and malate. As shown in Fig.4D-E and Fig. S4E neither state II nor state III respiration rates were altered with aging. However, physically active individuals displayed higher respiration rates supported by palmitoyl-L-carnitine and malate compared to their inactive counterparts (Fig.4D-E and Fig. S4E). Neither aging nor physical activity status impacted the ACR with P+M substrates (Fig. S4F).

To assess whether the greater respiration rates seen in active individuals resulted from a higher mitochondrial content, we assessed citrate synthase activity (CS), a robust marker of mitochondrial content^33,34^. As shown in Figures 4F and G, no effect of aging on CS activity was observed. However, physically active individuals displayed greater CS activity *vs* their inactive counterparts (Fig. 4F). Similarly to data on maximal respiration, a trend for a decrease in CS activity was observed in inactive participants (Fig. 4G). As with data on respiration, CS activity was positively associated with the number of daily steps in inactive participants (Fig. S4G). Importantly, when maximal respiration rates were normalized to CS activity, no effect of aging or physical activity status was observed (Fig. S4H-I). Collectively, our results indicate that aging *per se* in humans is associated with minimal to no change in mitochondrial respiration and content. They also indicate that physical activity status is associated with greater mitochondrial respiratory capacity and content throughout the human lifespan.

Since multiple previous studies had reported associations between mitochondrial respiration and parameters related to muscle mass, function, and physical performance ^12,29,35^, we assessed whether such associations would be found in our cohort. As shown in Figure S5, when all participants were pooled together, mitochondrial respiration was positively associated with muscle CSA, muscle strength and physical function.

### The impact of aging and physical activity status on skeletal muscle mitochondrial ROS production

Mitochondrial H_2_O_2_ emission (a widely used surrogate of mitochondrial ROS emission) was assessed in permeabilized myofibers using the Amplex Ultra Red system (Fig. 5A). As illustrated in Figure 5B-E and S6A-D neither state II or state III H_2_O_2_ emission rates supported by complex I or complex I and complex II substrates were affected by aging. The maximal H_2_O_2_ emission rate measured after the addition of oligomycin (ATP synthase inhibitor) and antimycin A (complex III inhibitor), was not impacted by aging (Fig.5D-E). Importantly, in all conditions tested, physically active participants displayed higher rates of H_2_O_2_ emission compared to inactive participants (Fig. 5B-E). No impact of aging or physical activity status on state II and state III H_2_O_2_ emission rates supported by lipid substrates (palmitoyl-L-carnitine and malate) was observed (Fig.5F-G and Fig. S6E-F). To assess whether aging and/or physical activity status impact intrinsically mitochondrial H_2_O_2_emission rates, the free radical leak was quantified by normalizing H_2_O_2_ emission to oxygen consumption rates. As can been seen in Figure S7, no impact of aging or physical activity status on mitochondrial free radical leak generation was observed. No negative association was observed between mitochondrial H_2_O_2_ emission and indices of physical function, strength and muscle mass (Figures S8 and S9). Taken together, our results indicate that aging *per se* in humans is not associated with an increase in the rate of mitochondrial H_2_O_2_ emission. They also indicate that being physically active results in greater H_2_O_2_ emission rates, an effect likely explained by a higher mitochondrial content in active individuals. Lastly, they indicate that intrinsic alterations in mitochondrial H_2_O_2_ emission are unlikely to contribute to the aging-related loss of muscle mass and function in humans.

**Figure 5:**
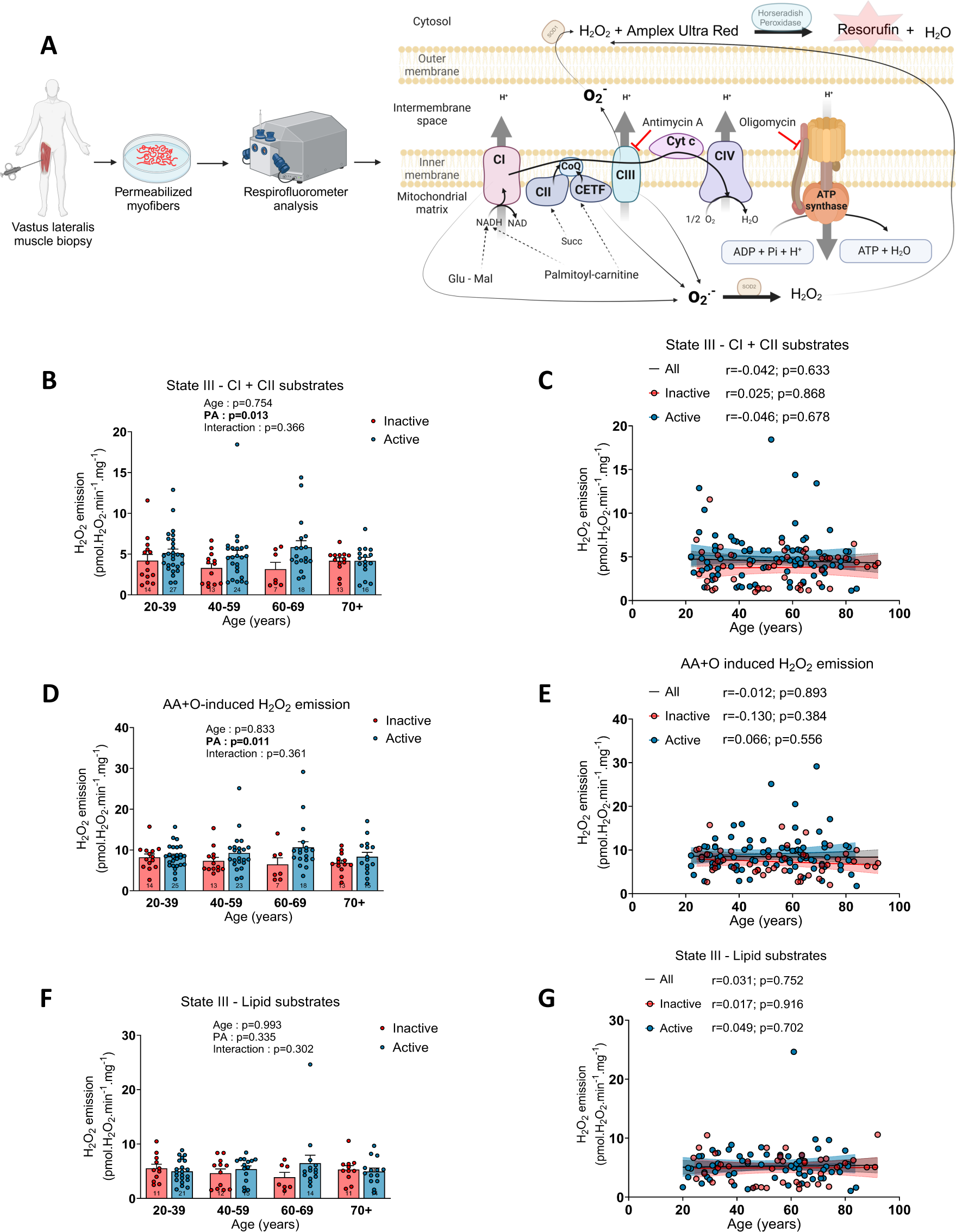
The impact of aging and physical activity status on skeletal muscle mitochondrial H2O2 emission. A) Schematic representation of the experimental design used to assess mitochondrialH2O2 emission in permeabilized myofibers. (B, C) State III H2O2 emission rates driven by complex I and II substrates (Glutamate + Malate + Succinate) in inactive and active participants. (D, E) Maximal H2O2 emission rates induced by the addition of Antimycin A (AA) and Oligomycin (O) in the presence of complex I and II substrates in inactive and active. (F, G) State III H2O2 emission rates driven by lipid substrates (palmitoyl-L-carnitine + malate) in inactive and active participants. For group analyses, results of the two-way ANOVA are displayed above each bar graph. Tukey post hoc tests were performed to test differences between age groups. Differences between inactive and active participants within each age group was assessed using multiple bilateral t-tests with FDR correction. Groups that do not share the same letter are significantly different. Linear regressions were performed to assess associations between age and variables of interest in the entire cohort (all) and for inactive and active participants separately. Pearson correlation coefficient (r) and p-values are displayed above each scatter plot. For two-way ANOVA, post hoc testing, and regression analyses, p<0.05 was considered statistically significant. For FDR analyses, q<0.05 was considered statistically significant. *=q<0.05. AA=antimycin A; O=oligomycin. Panel A was created with BioRender.com.

### The impact of aging and physical activity status on skeletal muscle mitochondrial calcium handling

The impact of aging and physical activity status on mitochondrial calcium handling remains an understudied aspect of mitochondrial biology in humans. In the present study, mitochondrial calcium handling was assessed in permeabilized phantom myofibers ^22,36^ by exposing mitochondria to a saturating calcium challenge. Such approach allows the measurement of the mitochondria calcium retention capacity (mCRC), of the time to mitochondrial permeability transition pore (mPTP) opening and of the mitochondrial calcium uptake rate (Fig.6A). The time to pore opening and mCRC were significantly reduced by aging (Fig. 6B-D and Fig. S10A and B). Post-hoc analyses for the mCRC showed that this reduction was significant between the 20-39 and 70+ and between the 60-69 and 70+ groups. Physical activity status had no impact on mCRC or time to pore opening (Fig.6B-C). As suggested by our ANOVA and post hoc analyses, further examinations of the relationship between aging and mCRC and the time to mPTP opening revealed that while both parameters were quite stable from 20 to 60 years of age, both sharply declined after 60 (Fig S10A-B). Interestingly, mCRC was correlated with markers of physical performance and muscle mass and function such as thigh lean mass, knee extension strength, and the performance at the 6 min walk test and step test (Fig.6E-H). A trend for a positive correlation between mCRC and muscle CSA was also observed when all participants were pooled together (Fig. S10C). No impact of aging on the mitochondrial calcium uptake rate was found in our cohort (Fig. S11). A trend for a higher calcium uptake rate in our physically active participants, mainly driven by participants aged 40 and over, was observed in our group analyses (Fig. S11A). Altogether, our data indicates that mitochondrial calcium handling is altered with aging independently from physical activity status and highlights positive associations between mCRC and muscle strength and physical performance.

**Figure 6:**
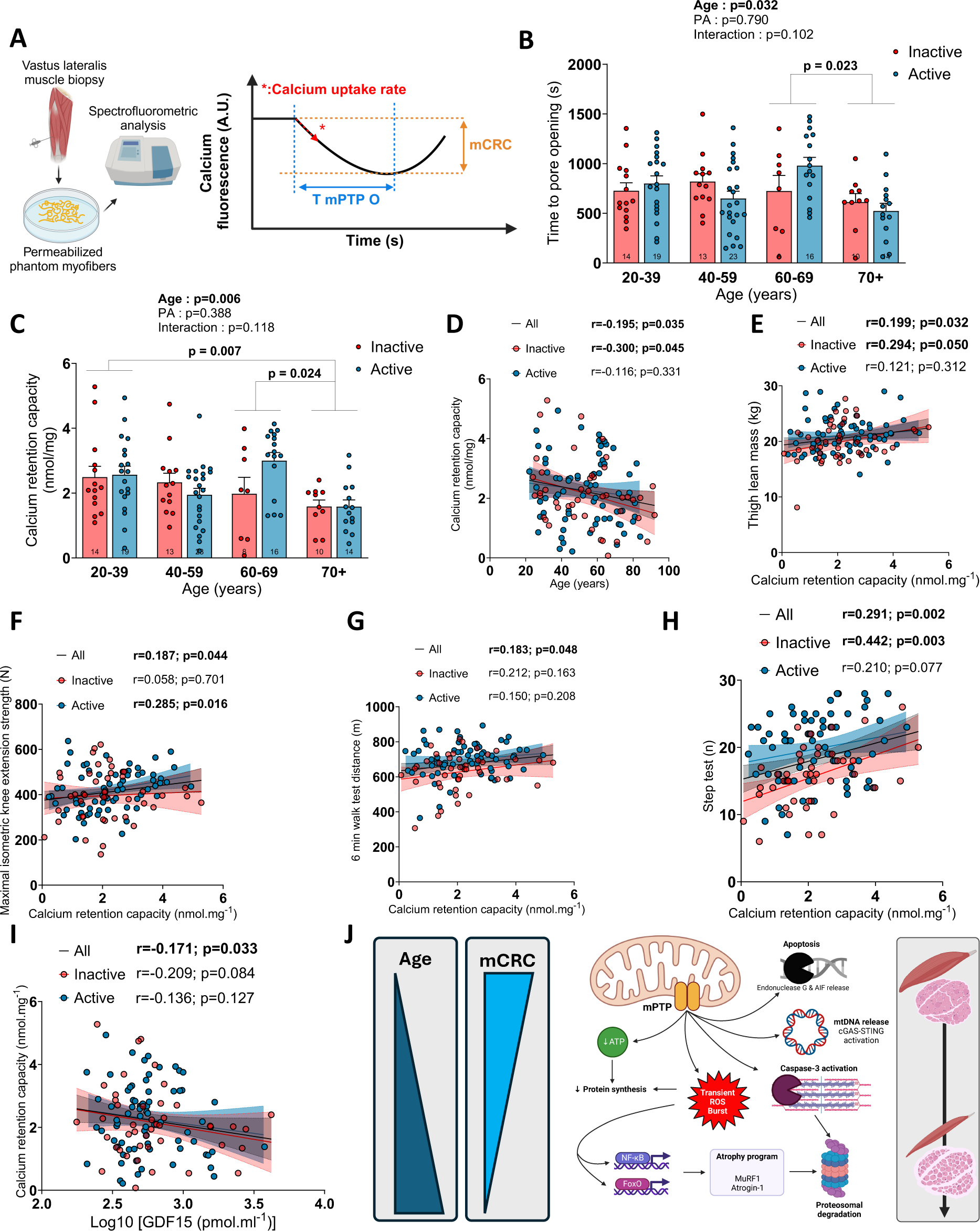
The impact of aging and physical activity status on skeletal muscle mitochondrial calcium handling. A) Schematic representation of the experimental design used to assess mitochondrial calcium retention capacity (mCRC), calcium uptake rate and time to mitochondrial permeability transition pore opening (T mPTP O) in permeabilized phantom myofibers. Quantification of the time to mitochondrial permeability transition pore opening (B) and mCRC (C,D) in inactive and active participants. Relationship between mCRC and (E) thigh lean mass, (F) maximal isokinetic knee extension strength, (G) distance at the 6 min walk test, (H) performance at the step test and (I) plasma GDF15 levels (log transformed) in our entire cohort (all) and for inactive and active participants separately. Three linear regressions are displayed in each graph to represent all, inactive and active participants only. (J) Schematic representation of the potential consequences of mPTP opening in muscle cells. For group analyses, results of the two-way ANOVA are displayed above each bar graph. Tukey post hoc tests were performed to test differences between age groups. Differences between inactive and active participants within each age group was assessed using multiple bilateral t-tests with FDR correction. Groups that do not share the same letter are significantly different. Linear regressions were performed to assess associations between age and variables of interest in the entire cohort (all) and for inactive and active participants separately. Pearson correlation coefficient (r) and p-values are displayed above each scatter plot. For two-way ANOVA, post hoc testing, and regression analyses, p<0.05 was considered statistically significant. For FDR analyses, q<0.05 was considered statistically significant. *=q<0.05. Panels A and J were created with BioRender.com.

### Association between various aspects of mitochondrial function and stress-responsive metabokine GDF15

Growth differentiation factor 15 (GDF15) has been described as a stress-responsive metabokine (metabolically-induced cytokine) and has recently emerged as potential key player in the human aging process (see ^37^ for a detailed review). Here we investigated whether specific aspects of mitochondrial biology in our cohort were associated with plasma GDF15 levels. As can be seen in Fig. S12A-B and Table 1, plasma GDF15 levels were exponentially and strongly correlated with age (r=0.83, p<0.0001) in the whole cohort and in active and inactive individuals separately. This confirms previous reports that have positioned GDF15 as a biomarker of human aging ^38–40^. Since the relationship between aging and plasma GDF15 levels was not linear (i.e. the aging-related increase in plasma GDF15 levels seemed to accelerate after 60), and because GDF15 values were not normally distributed on our cohort, log-transformed GDF15 values are used in regression analyses. Across the whole cohort, we find that individuals with higher plasma GDF15 levels had lower maximal mitochondrial respiratory capacity, a correlation which was stronger and only significant among inactive participants (Fig. S12C). No association between mitochondrial H_2_O_2_ emission and plasma GDF15 levels could be observed (Fig. S12D-F). However, individuals with higher levels of GDF15 had mitochondria more susceptible to undergo permeability transition (Fig. 6I). Taken altogether, these data relate skeletal muscle mitochondrial respiration and calcium handling with the systemic aging marker plasma GDF15.

## Discussion

While impairments in mitochondrial function are often portrayed as causally involved in the aging-related loss of muscle mass and function, evidence has emerged in the past few decades positioning physical activity as a major confounder ^23,24,41^. Until now, much of the research efforts have been focused on mitochondrial respiration and ROS production while other important mitochondrial functions, such as calcium handling, have been understudied in humans. To disentangle the relationship between aging, multiple aspects of mitochondrial function and physical activity in humans, we evaluated physical function, muscle mass and performance, muscle phenotype and various aspects of mitochondrial functions in 51 inactive and 88 active men between 20 to 93 years of age. Our data revealed that physical performance declines with age with partial protection from physical activity. While a trend for reduced maximal mitochondrial respiration was observed in inactive participants, no impact of aging was observed in active participants, indicating that aging *per se* does not alter mitochondrial respiration. Similarly, fatty acid (palmitoyl-carnitine) supported respiration was not altered by aging. Our data also indicate that physically active participants displayed higher fatty acid supported respiration as well as maximal respiration across the adult lifespan. Mitochondrial ROS production was unaffected by aging and physically active participants even displayed higher ROS production. Negative associations between mitochondrial ROS production and muscle mass, function or physical performance were not seen, indicating that mitochondrial ROS production is highly unlikely to play a role in the muscle aging process. In contrast, mitochondrial calcium retention capacity and time to mPTP pore opening decrease with aging regardless of the physical activity status. Mitochondrial calcium retention capacity also correlates with muscle strength and performance. Hence, from our data, we posit that alteration in mitochondrial calcium handling maybe a potential driver of muscle aging and that targeting mitochondrial calcium handling and/or the mitochondrial permeability transition pore may hold promise for treating age-related muscle impairments.

The present study lends strong credibility to the raising view that aging *per se* has limited impact on mitochondrial respiration ^14,22,25,27,42^. Indeed, while we observed a trend for lower mitochondrial respiration in physically inactive individuals, no decline in mitochondrial respiration could be observed in physically active participants (Figure 4C). Furthermore, maximal mitochondrial respiration was positively associated with the daily number of steps in inactive participants (Figure S4C) while the daily number of steps decreased with aging (Table 1), indicating that the trend for a lower maximal mitochondrial respiration with aging in inactive individual was consequential to reduced physical activity. Consistent with this view, it was shown that skeletal muscle mitochondria from older adults retain high plasticity in response to variation in mechanical load. Indeed, exercise training for relatively short terms (12 weeks) can substantially increase mitochondrial content ^43^ and respiration in older adults ^15,44,45^. Conversely, short term disuse (10 to 14 days of bedrest) results in a significant decrease in mitochondrial content ^46^ and respiration in older adults ^47,48^ while one hour of exercise per day can negate the effect of bed rest on mitochondrial bioenergetics in older adults ^48^. In contrast with previous reports suggesting that aging alters mitochondrial mass specific respiration (respiration normalized to a marker of mitochondrial content such as citrate synthase activity)^28^, our data show that normalizing respiration data to citrate synthase activity not only abolished the trend for lower maximal respiration with aging in inactive participants but also abolishes differences between active and inactive participants (Fig. S4E and I). These data indicate that maximal mitochondrial mass specific respiration is neither influenced by aging nor physical activity status. The absence of impact of physical activity status on maximal mitochondrial mass specific respiration is consistent with a recent report in human showing that differences in mitochondrial respiration seen between untrained, recreationally active and active-to-elite runners are abolished when normalized to mitochondrial volume or cristae density ^49^.

The absence of mitochondrial fatty acid oxidation change with age is also worth highlighting (Fig. 4D-E and Fig. S4H), especially when considering that fatty acids are important mitochondrial substrates at rest and during low intensity contractile activities, intensities most often used during activities of daily living. In contrast with previous studies that have reported data indicating mitochondrial uncoupling with muscle aging ^22,50,51^, the higher ACR values seen in the present and relatively large cohort using complex I substrates (Fig. S4D), and the absence of differences in ACR values when using lipids as substrates (Fig. S4F), indicate that muscle mitochondrial coupling efficiency is preserved with aging in humans. This view is in line with studies in rodents and humans that have reported no decrease in the ATP/O ratio (the most direct measure of mitochondrial coupling) in aged hindlimb skeletal muscles ^42,51,52^. Taken altogether, the data presented herein reinforce the necessity of considering physical activity when assessing muscle mitochondrial function, a requirement that likely extends to many others, if not all, aspects of muscle biology. Consideration related to physical activity (or lack of thereof) might also, at least in part, explain why mitochondrial respiration and content are often found impaired in old animals models of aging ^51–55^, albeit not always ^56^, as ambulatory activity was shown to decrease with aging in rodents ^57,58^. Importantly, and as reported in other studies ^12,29,35^, we also found in our cohort positive associations between mitochondrial respiration, muscle mass, strength, and power as well as physical performance (Figure S5). While our data indicate that a decrease in mitochondrial respiration is unlikely to drive the loss of muscle mass and function occurring during normal, healthy, aging, they nonetheless indicate that strategies boosting muscle mitochondrial function are likely to be beneficial to the health status of older adults. The association between maximal mitochondrial respiration and plasma GDF15 levels (a stress-responsive metabokine elevated in aging and aging-related diseases ^37^), particularly evident in inactive participants, further strengthen this view.

An increase in mitochondrial ROS production is often portrayed as a mechanism driving the muscle aging process ^16^. The data presented herein seriously challenge this view. Indeed, notwithstanding the use of many substrates and inhibitors of the mitochondrial oxidative phosphorylation, we did not see any effect of aging on mitochondrial H_2_O_2_ emission (Fig. 5 and S6), a widely used surrogate measure of mitochondrial ROS production ^22,42^. Even when H_2_O_2_ emission rates were normalized to the rate of oxygen consumption, a normalization that would reveal intrinsic alteration in ROS generation, no impact of aging was observed (Fig. S7). Physically active individuals, while clearly displaying greater physical performances from adulthood to old age, even displayed greater H_2_O_2_ emission compared to their inactive counterparts (Fig. 5 and S6). These data indicate that an increase in ROS production by the mitochondrial electron transfer system is highly unlikely to contribute to the aging-related loss of muscle mass and function in independently living men. Further strengthening this view, no negative association between mitochondrial H_2_O_2_ emission and muscle mass, function and physical performance was observed in the present study (Fig. S8 and S9). Our study, therefore, extends and support previous studies that have also reported that mitochondrial ROS production is unchanged with muscle aging ^22,42,56,59^. This view is also in line with previous reports that have found that long-term supplementation with the mitochondria-target antioxidants mitoquinone and SS-31 failed to rescue sarcopenia in rodents ^60,61^, while mice with MnSOD deficiency, an antioxidant enzyme located in the mitochondrial matrix, do not develop muscle atrophy with aging^62^.

The present study also details the impact of aging and physical activity on skeletal muscle composition. Indeed, the trend for an increase in the proportion of type I fibers in inactive individuals (Fig. 3D) is somewhat in line with data published by Lexell et al. over 3 decades ago^30^. Interestingly, such a trend was not observed in active participants and indicates a preservation of their fiber type II/I ratio. Consistent with previously published data ^27^, the analyses of MHC type IIa and type IIx positive myofibers (Fig. 3E-H) revealed that type IIa fibers seem particularly sensitive to aging as the proportion of pure type IIa fibers declined in both active and inactive participants. Combined with the trend for an increase in the proportion of type I fibers in our inactive group, this decrease in the proportion of type IIa fibers suggest a transition from type IIa to type I fibers in inactive individuals (Fig. 3). In contrast, the decrease in the proportion of type IIa fibers in active participants combined with an increase in the proportion of type IIa/IIx fibers indicate remodeling within the type II fibers category with aging in physically active individuals (Fig. 3). Based on data showing that exercise can improve the integrity of neuromuscular junctions (NMJ) in aged mice ^63^, it is tempting to speculate that these divergent phenotypic changes seen in physically active compared to physically inactive participants might reflect a protective impact of physical activity against alteration in NMJ in humans. While speculative at this stage, it should be noted that we employed an improved human muscle biopsy method to increase the probability of collecting NMJ ^64^. Immunohistological analyses on samples collected during the present study are underway and should allow us to get a better understanding of the effect of aging and physical activity on NMJ integrity in humans. The aging-related increase in intermuscular fat content is another important finding of the present study (Fig. 2H, I) as accumulating evidence position intermuscular adipose tissue as an important player in the aging-related deterioration of skeletal muscles ^65^. Our data also indicate that physical activity provides partial protection against this aging-related increase in intermuscular fat content (Fig. 2H, I).

Another important aspect of our work was to determine the impact of physical activity on mitochondrial calcium handling across the adult lifespan. By exposing permeabilized myofibers to a calcium challenge, we found that mitochondrial calcium retention capacity declines with aging independently of physical activity status (Fig. 6B, C). This finding is in line with previous studies that have reported a decline in mitochondrial calcium retention capacity with aging in recreationally active older adults ^22^ and even in master athletes ^66^. When mitochondrial calcium retention capacity is exceeded, opening of the mitochondrial permeability transition pore ensues. Our data support previous findings ^22^ indicating that the time to mitochondrial permeability transition pore opening was reduced with aging (Fig. 6C). These reduction in mCRC and time to mPTP opening demonstrate that muscle aging in humans is associated with mPTP sensitization (Fig. 6B-D). Our data indicate that the aging-related decline in mCRC and mPTP accelerates after 60 (Fig. S10A, B). Further highlighting the biological significance of these findings, we also report that mCRC is correlated with biologically and clinically relevant measures of muscle and physical function as well as with the stress responsive metabokine GDF15 (Fig. 6E-I). The fact that physical activity does not protect against the aging-related decline in mCRC and mPTP opening indicate that impaired calcium handling may be a mechanism driving muscle aging as even physically active individuals display aging-related muscle atrophy and weakness (Fig. 6). The aging-related alterations in calcium homeostasis that were reported in aged skeletal muscles, which include an increase in resting cytosolic calcium concentration ^67–70^ and reduced sarcoplasmic reticulum calcium uptake ^71^, are likely to synergize with a reduced mCRC and facilitate mPTP opening *in vivo*. Moreover, it was previously reported that the proportion of endonuclease G-positive myonuclei, a nuclease involved in apoptosis normally sequestered in the intermembrane space of mitochondria and released upon mPTP opening, increases with aging in both rodent and human skeletal muscles ^22,72^. Available data in rodents also provide strong support for an aging-related alteration of mCRC and mPTP function ^17,34,55^. Similarly, there is solid evidence that pro-apoptotic pathways downstream of mPTP opening are activated in aged rodent skeletal muscles ^53,72,73^. Since resistance training was shown to partially attenuate the aging-related decrease in sarcoplasmic reticulum calcium uptake ^71^, one could speculate that physical activity might maintain a cytosolic environment that decreases the probability of mPTP opening despite reduced mCRC in active individuals. Such preservation of the mitochondrial milieu might explain the partial protection conferred by physical activity on physical performance with aging. Several apoptosis-independent events could also link the aging-related sensitization of mPTP reported herein to altered muscle mass and function. Indeed, cytochrome C release upon mPTP opening leads to an increase in proteasomal activity secondary to caspase 3 activation ^74^. mPTP opening can also cause a transient burst in ROS production ^75^, which may stimulate the atrophy program through the activation of the FoxO transcription factor family ^76,77^. mPTP opening can also result in mitochondrial DNA release ^78,79^ and consequential activation of the cGAS-STING-NLRP3 inflammasome pathway, which was linked to upregulation of key ubiquitin ligases in the muscle atrophy program ^80,81^ (Figure 6J).

Collectively, the data presented in the present article indicate that intrinsic changes in mitochondrial respiration and ROS production are highly unlikely to drive the normal (non-pathological) muscle aging process. Our findings also strengthen the literature positioning physical activity as a powerful stimulus to improve mitochondrial energetics throughout the human adult lifespan. Furthermore, our results position altered mitochondrial calcium handling and sensitized mPTP as putative key events contributing to muscle atrophy and weakness that occurs in human aging (Figure 6J). Considering the presented data, a paradigm shift in a field overwhelmingly focused on mitochondrial bioenergetics may be needed. Finally, our results place mitochondrial calcium handling and the mPTP as potential therapeutic targets to treat the aging-related loss of muscle mass and function. Further studies will be required to test this exciting research avenue.

### Limitations of the study

Our study has some limitations. First, only male participants were enrolled. Due to the large sample size that the present study required and limited budget availability, we decided to focus our study on the sex with the most linear trajectory of neuromuscular aging ^82^. However, whether or not these findings are translatable to female participants remains to be established. Second, the present study enrolled only relatively healthy community-dwelling older adults as participants. Most older adults enrolled herein displayed relatively high physical performance, including those in the inactive group. In many ways, most of them matched the definition of successful aging ^83^. The magnitude and severity of mitochondrial changes with aging reported in the present study might therefore differ during states of pathological aging. Third, the cross-sectional design of our study is also limiting, preventing us from establishing a causal link between aging-related changes in mCRC and mPTP function with muscle weakness and reduced physical performance.

## Methods

### Participants

Participants were recruited by ads in newspaper, bulletin boards, social media and through word of mouth. To be included, participants had to be a man aged between 20 to 100 years old, with a BMI between 18 and 35 kg/m^2^. The following exclusion criteria were used: uncontrolled metabolic disease, neurodegenerative disease, smoking more than 2 cigarettes per day, drinking more than 2 glasses of alcohol per day, taking anticoagulant medication, having a pacemaker or a metal implant. Three separate on-site visits, organized within one month, were required to complete all measurement and procedures. All participants provided informed written consent after having received information on the nature, goal, procedures, and risks associated with the study. Participants were informed to maintain their physical activity and diets habits during their participation. To investigate the impact of aging and physical activity status on our variables of interest, participants were divided in the following age groups: young middle age (20-39 yo), mature middle age (40-59yo), young older adults (60-69 yo) and older adults (70+ yo) and sub-divided in inactive and active according to their physical activity status. All procedures were approved by the Ethics Committee of the *Université du Québec à Montréal* (UQÀM) (Ethic certificate number: 2020-2703). The study was conducted in the *Département des Sciences de l’activité physique* at UQÀM between February 2020 and June 2023.

### Physical activity assessment

To estimate the status of physical activity for each participant, we combined two approaches:

- Objective measures using a tri-axial accelerometer (SenseWear®Mini Armband) ^84^. Participants were asked to wear the armband on the non-dominant arm for at least 3 days and ideally for 7 days. Data were taken from days where the armband was worn at least 80% of the time over 24h (≥19h and 12 minutes). Data were analyzed using the sosware Armband Sensewear 8.1.
- Self-reported physical activity obtained through a structured interview with a trained kinesiologist. Questions asked during the interview were derived from the physical Activity Scale for the Elderly (PASE) ^85^. However, participants were asked to report their usual physical activity representative of the past 5 years and not just their physical activity over 7 days.

To be deemed active, participants had to fulfill at least one of the following criteria: i) Engage in a minimum of 150 minutes per week of structured moderate to intense physical activity, ii) Achieve a daily step count of at least 10,000 steps, or iii) Maintain a Metabolic Equivalent of Task (MET) equal to or greater than 1.6.

### Anthropometric data and body composition

Height (in cm) was measured using a wall-mounted height gauge (SECA 67029, Hanover, MD USA) and mass (in kg), with an electronic scale (ADAM - GFK 660a, USA). The BMI was calculated using the following formula: Mass (kg) / Height (m)^2^. DXA (GE Prodigy Lunar) was used to assess fat and lean masses. Participants were fasted for a minimum of two hours before the DXA scan.

### Assessment of muscle composition using p-QCT

Muscle composition was assessed using a peripheral quantitative computed tomography scan (p-QCT; Stratec XCT3000 STRATEC Medizintechnik GmbH, Pforzheim, Germany, Division of Orthometrix; White Plains, NY, USA). Images were taken in the lower third of the right thigh (from the right lateral epicondyle to the greater trochanter of the right femur). Acquisition parameters were the following: voxel size: 0.5 mm; speed :10 mm/s. p-QCT images were analyzed using ImageJ software (NIH, Bethesda, Maryland, USA, https://imagej.nih.gov/ij/) with the plugin p-QCT.

### Assessment of physical performance

Four validated tests were performed to assess physical performance as previously described Marcangeli *et al.* ^43^:

- 6 min walking test ^86,87^. The test was performed on a 25-meter track and participants were asked to walk as much as possible during 6 min. No encouragement was received along the test. The total distance was recorded in meters.
- alternative step test ^88,89^. Participants were placed facing toward a 20 cm height step and instructed to touch the top with the right and les foot, alternatively, as fast as possible during a 20 s period. The number of steps was recorded.
- “Sit to stand” test. Participants were asked to repeat standing up from a siwng position and siwng down as fast as possible for 30s with arms across their chests ^90^. The pme to perform 10 repeppons (10rep sit to stand) and the number of repetitions performed during 30s were recorded.
- “Timed Up & Go” test ^91^. This test, which consists in standing from a chair, walking a 3 m distance and siwng down again ^92^, was performed at a fast-paced walking speed. The time to perform this task was recorded.

### Assessment of muscle strength and power

The maximal isometric knee extension strength and the maximal lower limb muscle power were evaluated as previously described ^43^. Maximal isometric knee extension strength was assessed on the right leg using a strain gauge system attached to a chair (Primus RS Chair, BTE) upon which participants were seated with the knee and hip joint angles set at 135◦ and 90◦, respectively. The knee angle was set to 135◦, compared to the typical 90◦ to diminish the maximal joint torque that could be generated, particularly in light of generally more fragile bones in older adults. The tested leg was fixed to the lever arm at the level of the lateral malleoli on an analog strain gauge to measure strength. The highest of three maximum voluntary contractions was recorded.

Lower limb muscle power was measured on the right leg using the Nottingham Leg Extensor Power rig in a sitting position. Participants were asked to push the pedal down as hard and fast as possible, accelerating a flywheel attached to an analog to digital converter. Power was recorded for each push until a plateau/decrease was observed.

### Skeletal muscle biopsies

Muscles biopsies were collected using the Biopsy Electrostimulation for Enhanced NeuroMuscular Junction Sampling (BeeNMJs) method as extensively described ^64^. This approach, which allows for enrichment of NMJ in muscle biopsy samples, was selected as our overall research project was aimed at investigating the contributions of mitochondrial dysfunction and altered NMJ integrity in the human muscle aging process. Analyses of NMJ integrity are currently underway and will be part of a separate manuscript. The first step of the BeeNMJs method consist in identifying the region of the VL with the highest NMJ density through electrostimulation ^64^. Once identified, muscle biopsies were performed along the main axis of VL at least 2 cm away from the site with the highest NMJ density. Muscle biopsies were performed using a suction modified Bergström needle under local anesthesia (lidocaine injection). The tissue collected during the biopsy procedures was either used fresh to assess mitochondrial function, prepared for histological analyses, prepared to assess NMJ integrity or snap frozen in liquid nitrogen and stored at -80°C until use.

### Preparation of permeabilized muscle fibers for *in situ* assessment of mitochondrial function

Mitochondrial function was assessed in freshly muscle biopsy samples. Once dissected, muscle biopsy samples were weighted with a precision scale and then rapidly immersed in ice-cold stabilizing buffer A (2.77 mM CaK_2_ ethylene glycol-bis-(2-aminoethylether)-N,N,N,N-tetraacetic acid (EGTA), 7.23 mM K_2_ EGTA, 6.56 mM MgCl_2_, 0.5 mM dithiothreitol (DTT), 50 mM 2-(N-morpholino) ethanesulfonic acid potassium salt (K-MES), 20 mM imidazol, 20 mM taurine, 5.3 mM Na_2_ ATP, and 15 mM phosphocreatine, pH 7.3). Muscle biopsy samples were separated into small fiber bundles using fine forceps under a surgical dissecting microscope (Leica S4 E, Germany). Muscle fiber bundles were incubated into a glass scintillation vial for 30 minutes at low rocking speed containing buffer A supplemented with 0.05 mg/mL saponin (Sigma) to selectively permeabilize the sarcolemma. Fiber bundles was divided in two parts, once was washed 3 times 10 minutes at low rocking speed in the MiR05 buffer (110 mM sucrose, 20 mM HEPES, 10 mM KH_2_PO_4_, 20 mM taurine, 60 mM K-lactobionate, 3 mM MgCl_2_, 0.5 mM EGTA, 1 g/L of fatty acid free BSA, pH 7.4) to assess mitochondrial respiration and H_2_O_2_ emission. The other part of fiber bundles was washed 3 times 10 minutes at low rocking speed in the C buffer (80mM K-MES, 50mM HEPES, 20mM taurine, 0.5mM DTT, 10mM MgCl_2_, 10mM ATP, pH 7.3) to realize mPTP analyses.

### Assessment of mitochondrial respiration

Permeabilized myofibers from muscle biopsy samples respiration was performed using an Oroboros O2K high-resolution fluororespirometer (Oroboros Instruments, Austria) at 37 °C in 2 mL of MiR05 buffer. Briefly, 3 to 6 mg (wet mass) of permeabilized fiber bundles were weighed and added to the respiration chambers. The following 2 protocols were used:

- Protocol 1 consisted in the sequential addition of the following substrates and inhibitors: 10 mM glutamate + 5 mM malate (G+M), 2 mM ADP, 10 mM succinate, 1µM oligomycin and 2µM antimycin A.
- Protocol 2 consisted in the sequential addition of the following substrates and inhibitors: 200µM mM palmitoyl-L-carnitine + 5 mM malate, 2 mM ADP.

After mitochondrial respiration measurements, bundles were removed and placed in liquid nitrogen and stored at -80°C for the assessment of citrate synthase (CS) activity. Respiration rates were normalized as nanomoles of dioxygen per minute per mg of wet muscle mass. All respiration experiments were analyzed with MitoFun ^93^, a homemade code to analyze mitochondrial function data in the Igor Pro 8 software (Wavemetrics, OR, USA).

### Assessment of mitochondrial H_2_O_2_ emission

The H_2_O_2_ emission from myofiber bundles was assessed by monitoring the rate of H_2_O_2_ release using the amplex ultra red-horseradish peroxidase system. This was performed along with respiration assessment in the Oroboros O2K high-resolution fluororespirometer (Oroboros Instruments, Austria) at 37 °C in 2 ml of MiR05 buffer supplemented with Amplex Ultra Red (10μM), SOD (5 U/ml), and HRP (1 U/ml). A calibration curve was generated daily using successive additions of known [H_2_O_2_] in absence of tissue. After mitochondrial H_2_O_2_ emission measurements were completed, bundles were retrieved, snap frozen in liquid nitrogen and stored at -80°C until CS activity measures. H_2_O_2_ emission was normalized as picomoles per minute per milligram of wet muscle mass. All H_2_O_2_ emission experiments were analyzed with MitoFun ^93^, a homemade code to analyze mitochondrial function data in the Igor Pro 8 software (Wavemetrics, OR, USA).

### Assessment of mPTP sensitivity to Calcium

Sensitivity to mPTP opening was assessed by determining mitochondrial CRC in the presence of a calcium challenge ^36^. The high affinity for Ca^2+^ of myosin-actin-associated proteins prevents measurements of mitochondrial Ca^2+^ uptake in standard permeabilized myofibers. To overcome this limitation, phantom fibers (fibers without myosin) were prepared ^36^. As mentioned earlier, fiber bundles were first permeabilized with saponin and washed 3 times in buffer C. The bundles were then incubated for 30 min with intermittent manual agitation at 4°C in buffer D (800 mM KCl, 50 mM HEPES, 20 mM taurine, 0.5 mM DTT, 10 mM MgCl_2_, and 10 mM ATP, pH 7.3 at 4°C), a solution of high ionic force to extract myosin but which preserves mitochondrial function (phantom fibers). Bundles were finally washed 3 times in low-EGTA CRC buffer (250 mM sucrose, 10 mM Tris, 5 μM EGTA, and 10 mM 3-(*N*-morpholino) propane sulfonic acid (MOPS), pH 7.3 at 4°C) and kept on ice until use for Ca^2+^-induced mPTP opening assays. The CRC was then determined in permeabilized phantom fibers as previously described ^22^. Briefly, a muscle bundle of 3–6 mg wet mass was added to 600 μl of CRC buffer containing ∼30 μM of Ca^2+^ supplemented with 5 mM glutamate, 2.5 mM malate, 10 mM inorganic phosphate (P_i_), 1 μM calcium-green 5 N, and 0.5 µM oligomycin. Mitochondrial Ca^2+^ uptake was immediately followed by monitoring the decrease in extramitochondrial Ca^2+^concentration using the fluorescent probe calcium-green 5 N (C3737, ThermoFisher). Fluorescence was detected using a spectrophotometer (Hitachi F2500, FL Solutions software) with excitation and emission detectors set at 505 and 535 nm, respectively. All measurements were performed at 37°C. Progressive uptake of Ca^2+^ by mitochondria was monitored until mitochondrial Ca^2+^ release caused by opening of the mPTP was observed as the inversion of signal. CRC was calculated as total amount of Ca^2+^ taken by mitochondria before Ca^2+^ release. Fluorescent units were converted into Ca^2+^ concentration using a standard curve obtained daily using successive addition of known Ca^2+^ concentration in the absence of sample. CRC values were expressed per milligram of wet fiber mass. The time pore to opening was also calculated as a time that mitochondria take to open the transition pore and release the Ca^2+^ and data were expressed in second. All mPTP experiments were analyzed with MitoFun ^93^, a homemade code to analyze mitochondrial function data in the Igor Pro 8 software (Wavemetrics, OR, USA).

### Assessment of citrate synthase activity

Enzymatic activity level of the mitochondrial citrate synthase (CS) was determined spectrophotometrically on permeabilized myofibers frozen immediately after respirometry and H_2_O_2_ emission assays. Briefly, permeabilized myofibers were homogenized in 20 volumes of an extraction buffer containing 1mM EDTA, Triethanolamine 50mM. The homogenates were centrifuged at 10,000g for 2 min at 4°C and incubated on ice for 1h. The supernatants were collected in a clean Eppendorf and CS activity was measured spectrophotometrically by detecting the increase in absorbance at 412 nm in a 96-well plate, at 30°C, using 200μL of a reaction buffer (200 mM Tris, pH 7.4) containing 2µM acetyl-CoA, 200µM 5,5ʹ-dithiobis-(2-nitrobenzoic acid) (DTNB), 350 µM oxaloacetic acid, and 0.1% Triton-X100. The kinetic assay was performed with the M1000 TECAN multiplate reader for 2 min with repeated measures every 5 seconds. To calculate the CS activity, the DTNB molar extinction coefficient used was 13.6 L. mol^−1^.cm^−1^. CS activity absorbance units were normalized to mg of frozen fibers.

### Histology analyses

#### Skeletal muscle sample sectioning

Muscle biopsies samples were mounted in tragacanth (Sigma, G1128), frozen in cooled liquid isopentane, and stored at -80°C. Ten-micron thick serial cross-sections were cut in a cryostat at -20°C and mounted on lysine coated slides (Superfrost).

#### Assessment of muscle fiber type and size

Muscle cross-sections were immunolabelled for different MHC isoforms as previously described ^27,94^. Briefly, muscle cross sections were allowed to reach room temperature, rehydrated with Phosphate Buffer solution (PBS, Sigma, 4417) and blocked with PBS containing 10% of goat serum (GS) for 45 min at room temperature. Muscle sections were then incubated for 1 hour, at room temperature with the following primary antibodies mix: a mouse IgG2b monoclonal anti-MHC type I (BA-F8, 1:25, DSHB, Iowa, IA, USA), a mouse IgG1 monoclonal anti-MHC type IIa (Sc-71, 1:200, DSHB, Iowa, IA, USA), a mouse IgGM monoclonal anti-MHC type IIx (6-H1, 1:25, DSHB, Iowa, IA, USA) and a rabbit IgG polyclonal anti-laminin (Sigma, L9393, 1:750) diluted in PBS containing 10% of GS. Muscle cross-sections were then washed 3 times with PBS before being incubated for 1 hour with the following secondary antibodies mix diluted in PBS containing 10% of GS: Alexa Fluor 350 IgG2b goat anti-mouse (A21140), Alexa Fluor 594 IgG1 goat anti-mouse (A21125), Alexa Fluor 488 IgM goat anti-mouse (A21042) and Alexa Fluor 488 IgG goat anti-rabbit (A11008). Sections were washed 3 times in PBS and cover slipped using Prolong Gold (ThermoFisher, P36934) as mounting medium. Slides were imaged using an Olympus IX83 Ultra Sonic fluorescence microscope (Olympus, Japan). Analyses of fiber type and size were performed by an experimenter blinded to the participants age and physical activity status. At least 200 fibers were manually traced per participants (average ± SD: 314 ± 86) using the ImageJ software (NIH, Bethesda, Maryland, USA, https://imagej.nih.gov/ij/).

#### General blood biochemistry

Fifteen ml of of veinous blood sample were collected after an overnight fast in BD Vacutainer SST Tube with Hemogard Closure (Gold, Dufort et Lavigne Ltée, Qc, Canada, BEC367986). Tubes were inverted 6 times and were then left at room temperature for a minimum of 30 min and maximum of 60 minutes. Samples were then stored at 4°C, for a maximum of 4 hours, before centrifugation. Tubes were centrifuged at 2000g for 15 minutes at room temperature. After centrifugation, the supernatant was collected and aliquoted into 1.5 ml Eppendorf tubes and immediately stored at -80°C until use.

Serum glucose concentrations were assessed using a coupled enzymatic assay in a Beckman Coulter AU analyzer according to the manufacturer instructions (Beckman Coulter, BAOSR6X21). Serum insulin concentrations were assessed using the ARCHITECT Insulin assay according to the manufacturer instructions (Abbott, ARCHITECT Insulin 8K41). Fructosamine levels were assessed using a colorimetric assay according to the manufacturer instruction (Randox, FR 3133).Total cholesterol and HDL-cholesterol were assessed using a Beckman Coulter AU analyzer according to the manufacturer instructions (Beckman Coulter, BAOSR6X16 and BAOSR6x95, respectively). LDL-cholesterol was calculated based on HDL-cholesterol and triglycerides values as follows: LDL cholesterol = total cholesterol - HDL - (triglycerides / 5). The HOMA-IR was calculated as [fasting glucose (mmol/l) x fasting insulin (pmol/l)]/135 ^95^. The Quantitative Insulin Sensitivity Check Index (QUICKI) 1 ⁄ [log fasting insulin (μU⁄mL) + log glucose (mg⁄dL)] ^96^. CRP levels were assessed using the Beckman Coulter CRP latex assay in a Beckman Coulter AU analyzer according to the manufacturer instructions (Beckman Coulter, BAOSR6X99). Abnormally high CRP values for two participants (one in the 60-69 inactive group and one in the 70+ active group) were removed from the analyses. Triglycerides levels were assessed using a colorimetric assay in a Beckman Coulter AU analyzer according to the manufacturer instructions (Beckman Coulter, BAOSR6X118). Serum protein levels were assessed using a colorimetric assay in a Beckman Coulter AU analyzer according to the manufacturer instructions (Beckman Coulter,BAOSR6X32).

#### Quantification of circulating GDF15 level

Fifteen ml of veinous blood sample were collected after an overnight fast in tubes containing ethylenediaminetetraacetic acid (EDTA). Tubes were inverted 10 times and then stored at 4°C, for a maximum of 4 hours, before centrifugation. Tubes were centrifuged at 2000g for 15 minutes at room temperature. After centrifugation, the supernatant was collected and aliquoted into 1.5 ml Eppendorf tubes and stored at -80°C until use. Plasma GDF15 levels were quantified using a high-sensitivity ELISA kit (R&D Systems, DGD150, SGD150) as described previously ^97^.

### Statistical analyses

All statistical analyses were performed using GraphPad Prism 10.1.3. Group comparisons were performed using two-way analysis of variance (ANOVA; p<0.05 were considered significant) with age and physical activity status as factors. When an age effect was observed, Tukey’s multiple comparisons tests were performed to assess differences between age groups (p<0.05 were considered significant). To assess whether differences were present between active and inactive participants in each age group, bilateral students t-tests followed by a correction using the false discovery rate (FDR) (using the two-stage step-up method of Benjamini, Krieger, and Yekutieli) was applied (p < 0.05 and q < 0.05 were considered significant). For analyses where age was considered as a continuous variable, and for all correlation analyses, simple linear regressions were performed, and Pearson r coefficients were calculated. To test whether correlations were significant, two-tailed tests were used were used for all correlations except for correlations involving plasma GDF15 levels which were tested with one-tailed tests. Bar graphs are presented as mean ± standard error of the mean (SEM). The exact number of participants within each group or analyses in all figures is indicated either in bar graphs or in the figure legends. Individual data are displayed in each graph.

## Acknowledgements

We are grateful to all study participants who donated their time and tissues. We sincerely thank Monica Tanase and Manon Dargegen for their technical assistance during muscle biopsies and for histological analyses. We thank Didier Brassard for insightful discussions on statistical analyses. We are grateful to the CERMO-FC for granting us access to its Cellular and imaging analyzes and Biophysics and biomolecular screening platforms.

## Author contributions

M.C., V.M., P.G., M.A.-L., M.B., R.R., J.A.M. and G.G. designed the research. P.G., M.A.-L., M.B., R.R., J.A.M. and G.G secured funding. M.C., V.M., R.H., J.G., J.-P.L.-G., J.A.M., C.T., H.H., M.P. and G.G. performed the experiments. All authors contributed to data analysis and interpretation. M.C., V.M. and G.G. wrote the original draft with inputs from all authors. All authors read, edited, and approved the manuscript.

