## Supplementary Figures for "The impact of physical activity on physical performance, mitochondrial bioenergetics, ROS production and calcium handling across the human adult lifespan"

**Figure S1:** Supplementary data on the impacts of aging and physical activity status on physical function, muscle strength and power.

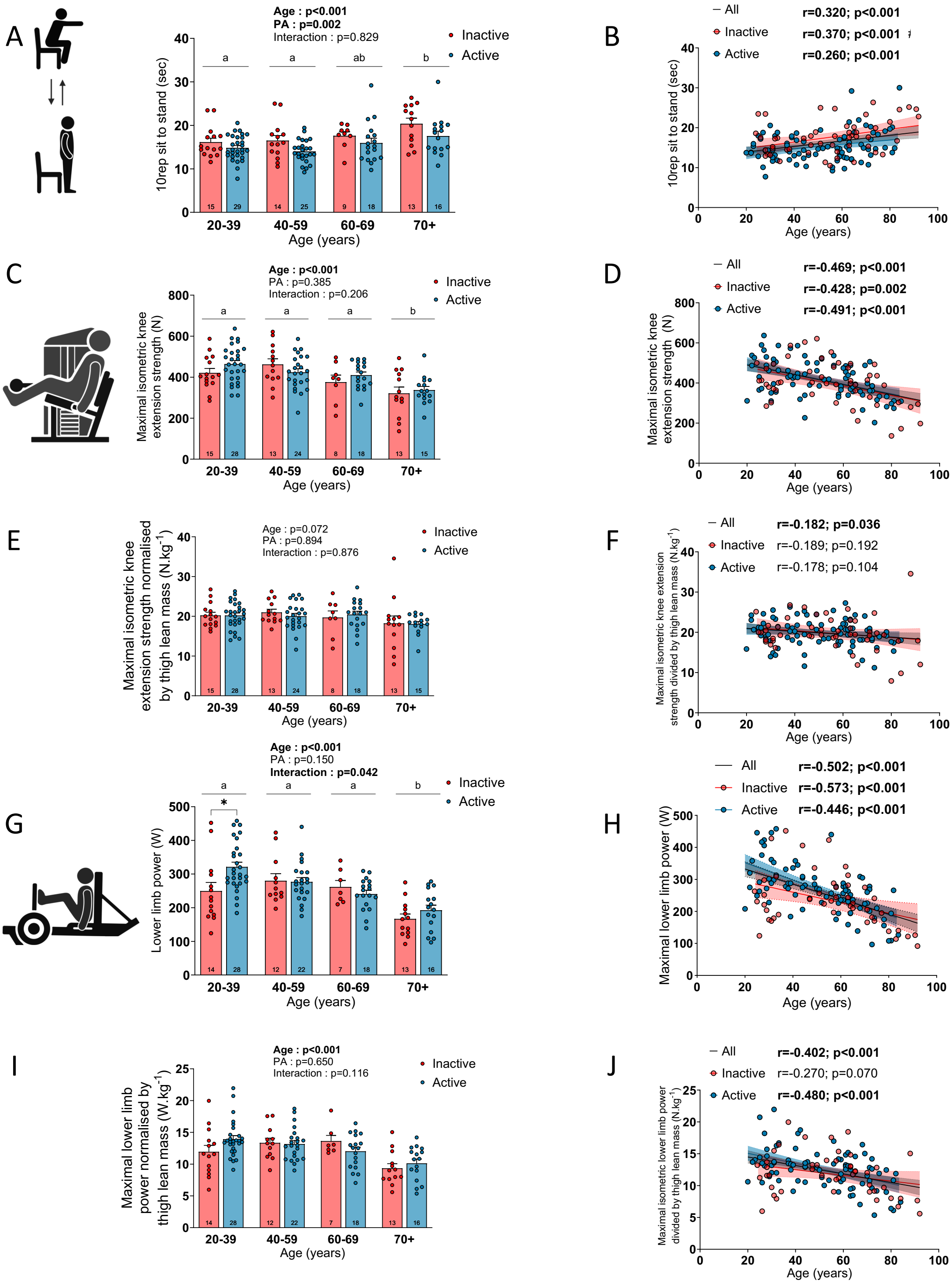


**Figure S1: Supplementary data on the impacts of aging and physical activity status on physical function, muscle strength and power.**

Quantification of the performance at the (A-B) sit to stand test (time to perform 10 repetitions) in inactive and active participants. Quantification of the maximal isometric knee extension strength (C-D), maximal isometric knee extension strength normalized to thigh lean mass (E-F), maximal lower limb power (G-H) and maximal lower limb power (I-J) normalized to thigh lean mass in inactive and active participants. For group analyses, results of the two-way ANOVA are displayed above each bar graph. Tukey post hoc tests were performed to test differences between age groups. Differences between inactive and active participants within each age group was assessed using multiple bilateral t-tests with FDR correction. Groups that do not share the same letter are significantly different. Linear regressions were performed to assess associations between age and variables of interest in the entire cohort (all) and for inactive and active participants separately. Pearson correlation coefficient (r) and p-values are displayed above each scatter plot. For two-way ANOVA, post hoc testing, and regression analyses, $p<0.05$ was considered statistically significant. For FDR analyses, $q<0.05$ was considered statistically significant. *=$q<0.05$. Icons in panels A, C and G were created with BioRender.com.

**Figure S2:** The impacts of aging and physical activity status on thigh lean mass and subcutaneous fat content.

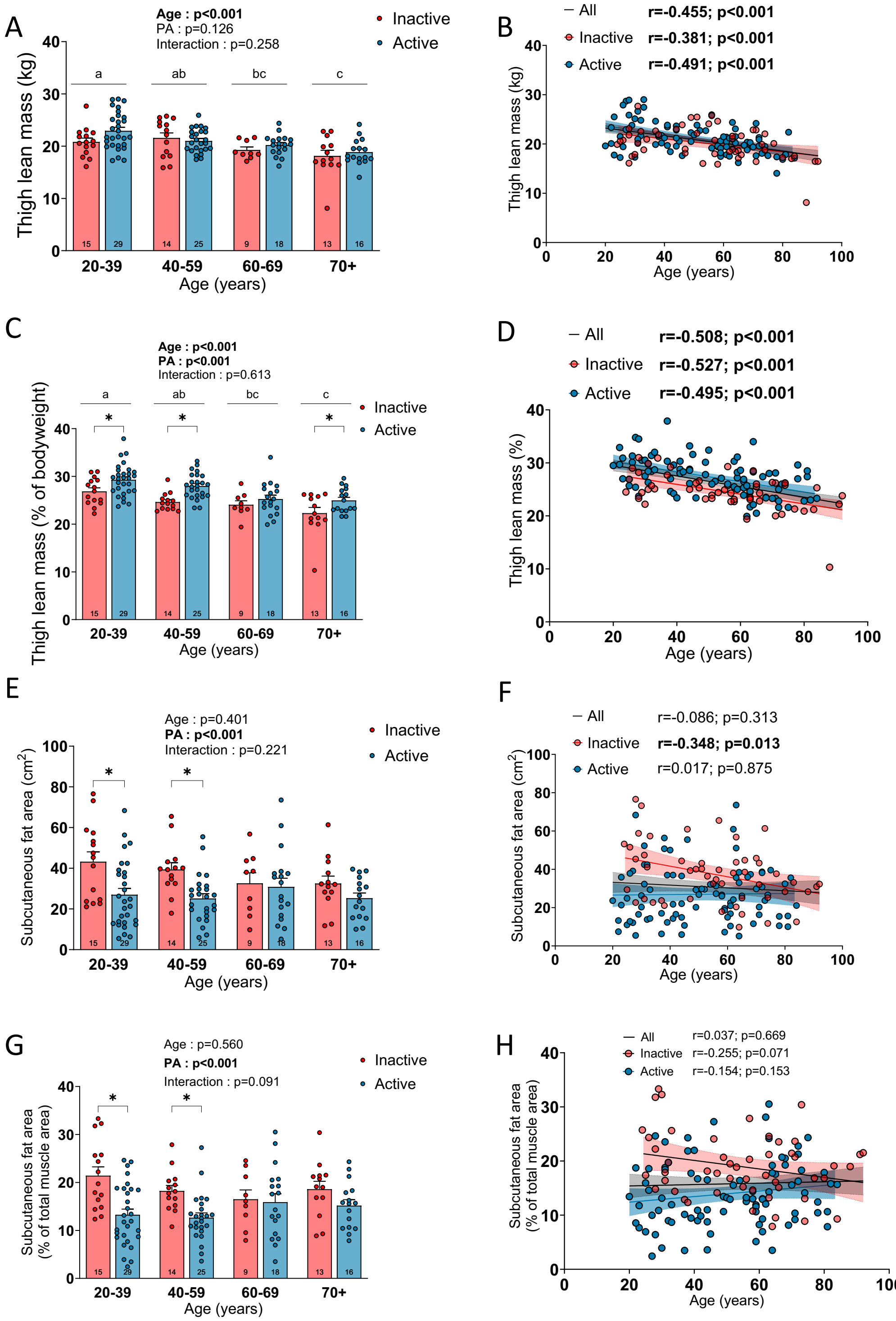


**Figure S2: The impacts of aging and physical activity status on thigh lean mass and subcutaneous fat content.**

Quantification of the thigh lean mass (A-B) and thigh lean mass normalized to total body weight (C-D) in inactive and active participants. Quantification of the subcutaneous fat area

(E-F) in inactive and active participants. For group analyses, results of the two-way ANOVA are displayed above each bar graph. Tukey post hoc tests were performed to test differences between age groups. Differences between inactive and active participants within each age group was assessed using multiple bilateral t-tests with FDR correction. Groups that do not share the same letter are significantly different. Linear regressions were performed to assess associations between age and variables of interest in the entire cohort (all) and for inactive and active participants separately. Pearson correlation coefficient (r) and p-values are displayed above each scatter plot. For two-way ANOVA, post hoc testing, and regression analyses, p<0.05 was considered statistically significant. For FDR analyses, q<0.05 was considered statistically significant. *=q<0.05.

**Figure S3:** Supplementary data on the impacts of aging and physical activity status on muscle area and fiber type proportion.

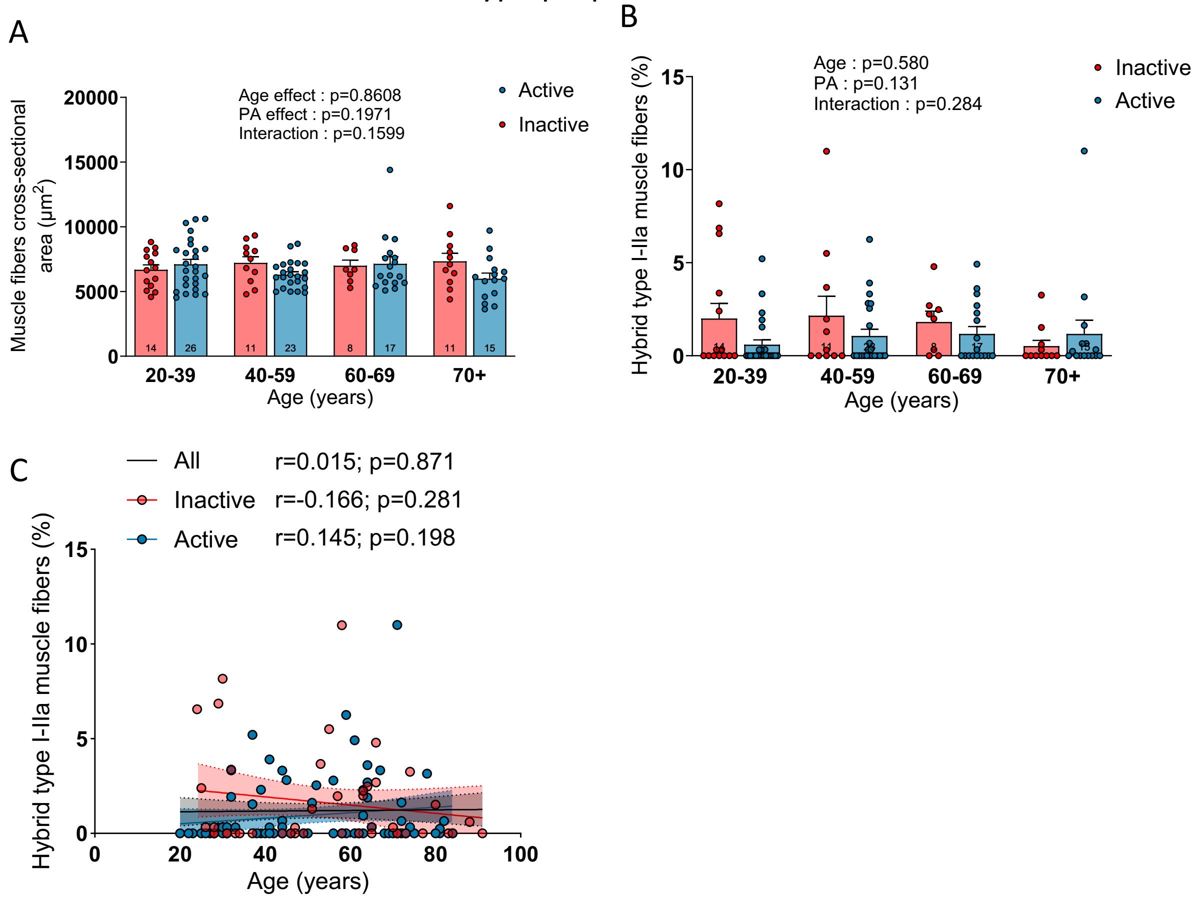


**Figure S3: Supplementary data on the impacts of aging and physical activity status on muscle area and fiber type proportion.**

Quantification of the muscle fibers cross-sectional area (A) and the proportion of hybrid type I-IIa muscle fibers (B-C). For group analyses, results of the two-way ANOVA are displayed above each bar graph. Tukey post hoc tests were performed to test differences between age groups. Differences between inactive and active participants within each age group was assessed using multiple bilateral t-tests with FDR correction. Groups that do not share the same letter are significantly different. Linear regressions were performed to assess associations between age and variables of interest in the entire cohort (all) and for inactive and active participants separately. Pearson correlation coefficient (r) and p-values are displayed above each scatter plot. For two-way ANOVA, post hoc testing, and regression analyses, $p<0.05$ was considered statistically significant. For FDR analyses, $q<0.05$ was considered statistically significant.

**Figure S4:** Supplementary data on the impacts of aging and physical activity status on mitochondrial respiration.

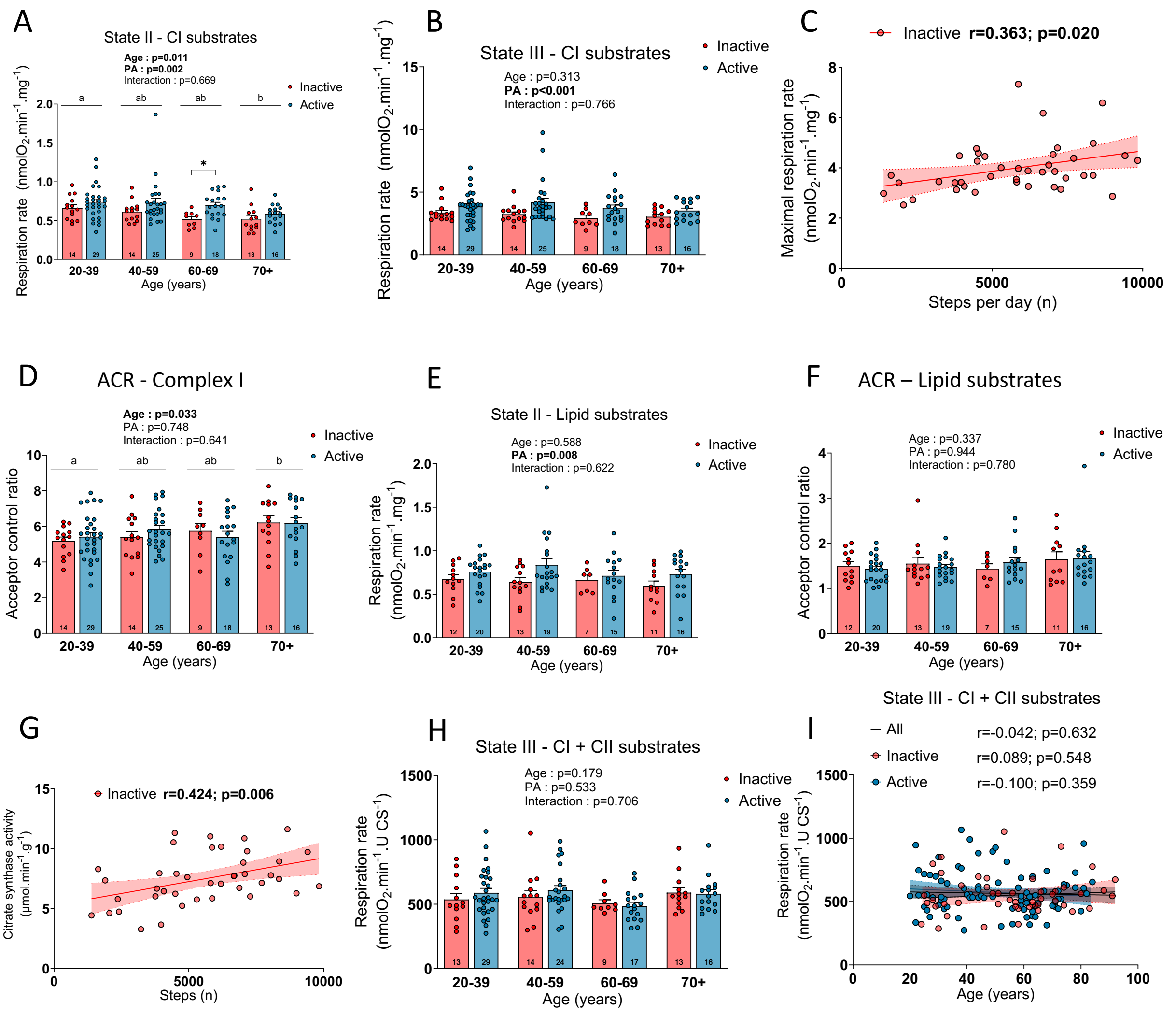


**Figure S4: Supplementary data on the impacts of aging and physical activity status on mitochondrial respiration.**

(A) State II and (B) state III (ADP stimulated) respiration rate driven by complex I substrates (Glutamate + Malate) in inactive and active participants. (C) Correlations between the state III (ADP stimulated) respiration rate with complex I (Glutamate + Malate) and II (Succinate) substrates and number of steps per day in inactive participants. (D) Acceptor control ratio, calculated by dividing state III by state II respiration rates with glutamate and malate substrates, in inactive and active participants. (E) State II respiration rate driven by lipids substrates (Palmitoyl-L-carnitine and Malate) in inactive and active participants. (F) Acceptor control ratio, calculated by dividing state III by state II respiration rates with lipids substrates (Palmitoyl-L-carnitine and Malate), in inactive and active participants. (G) Correlations between citrate synthase activity and number of steps per day in inactive participants. (H-I) state III (ADP stimulated) respiration rate driven by complex I (Glutamate + Malate) and II (Succinate) substrates normalized by citrate synthase activity in inactive and active participants. For group analyses, results of the two-way ANOVA are displayed above each bar graph. Tukey post hoc tests were performed to test differences between age groups. Differences between inactive and active participants within each age group was assessed using multiple bilateral t-tests with FDR correction. Groups that do not share the same letter are significantly different. Linear regressions were performed to assess associations between age and variables of interest in the entire cohort (all) and for inactive and active participants separately. Pearson correlation coefficient (r) and p-values are displayed above each scatter plot. For two-way ANOVA, post hoc testing, and regression analyses, $p<0.05$ was considered statistically significant. For FDR analyses, $q<0.05$ was considered statistically significant. *=$q<0.05$.

**Figure S5:** Correlations between maximal mitochondrial respiration and physical function.

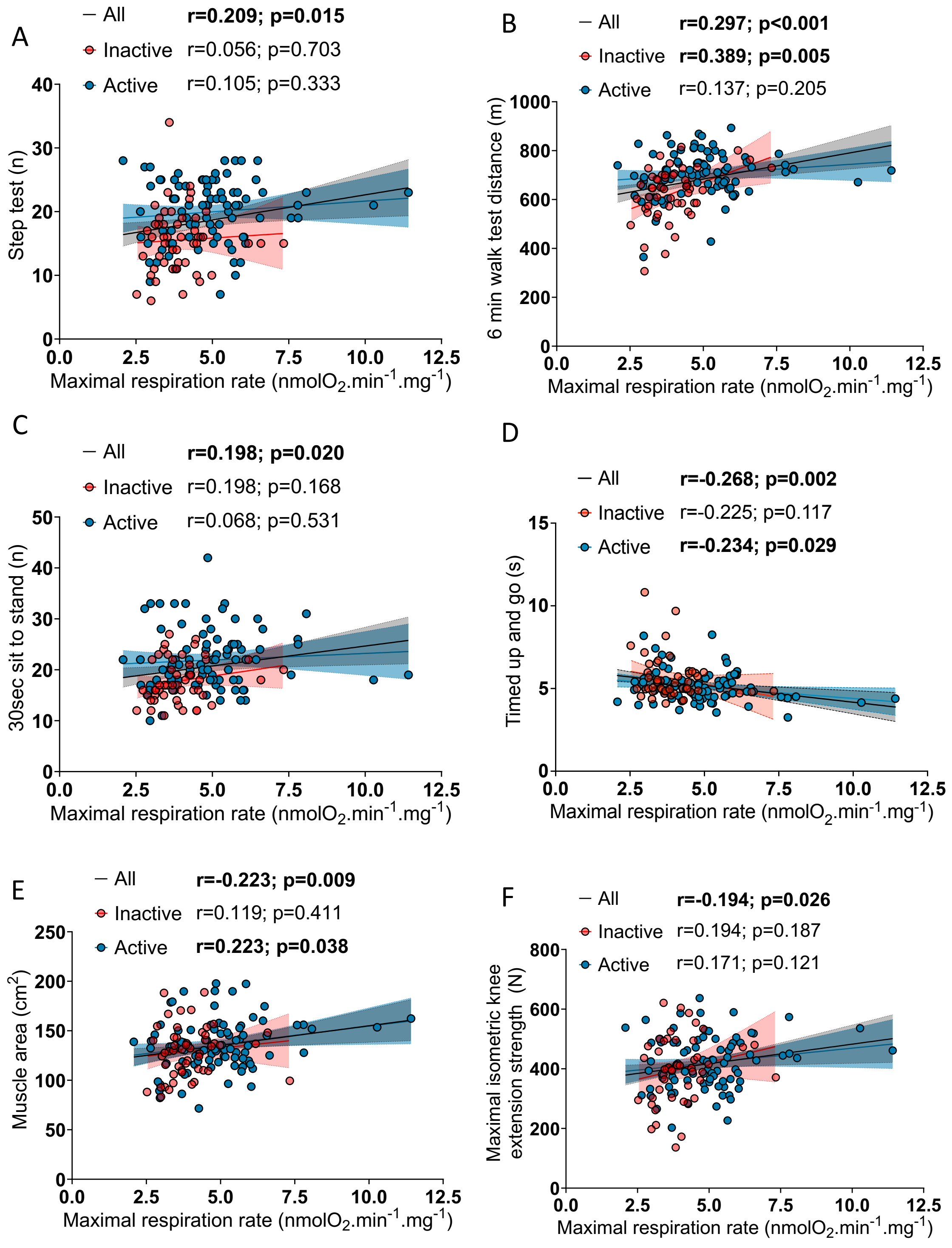


**Figure S5: Correlations between maximal mitochondrial respiration and physical function.** Correlations between state III (ADP stimulated) respiration rate driven by complex I (Glutamate + Malate) and II (Succinate) substrates and (A) step test, (B) 6 min walk test distance, (C) sit to stand test (number of repetitions performed in 30s), (D) timed up and go, (E) muscle cross-sectional area and (F) maximal isometric knee extension strength in inactive and active participants. Linear regressions were performed to assess associations between age and variables of interest in the entire cohort (all) and for inactive and active participants separately. Pearson correlation coefficient (r) and p-values are displayed above each scatter plot. p<0.05 was considered statistically significant.

**Figure S6:** Supplementary data on the impacts of aging and physical activity status on mitochondrial $H_2O_2$ emission.

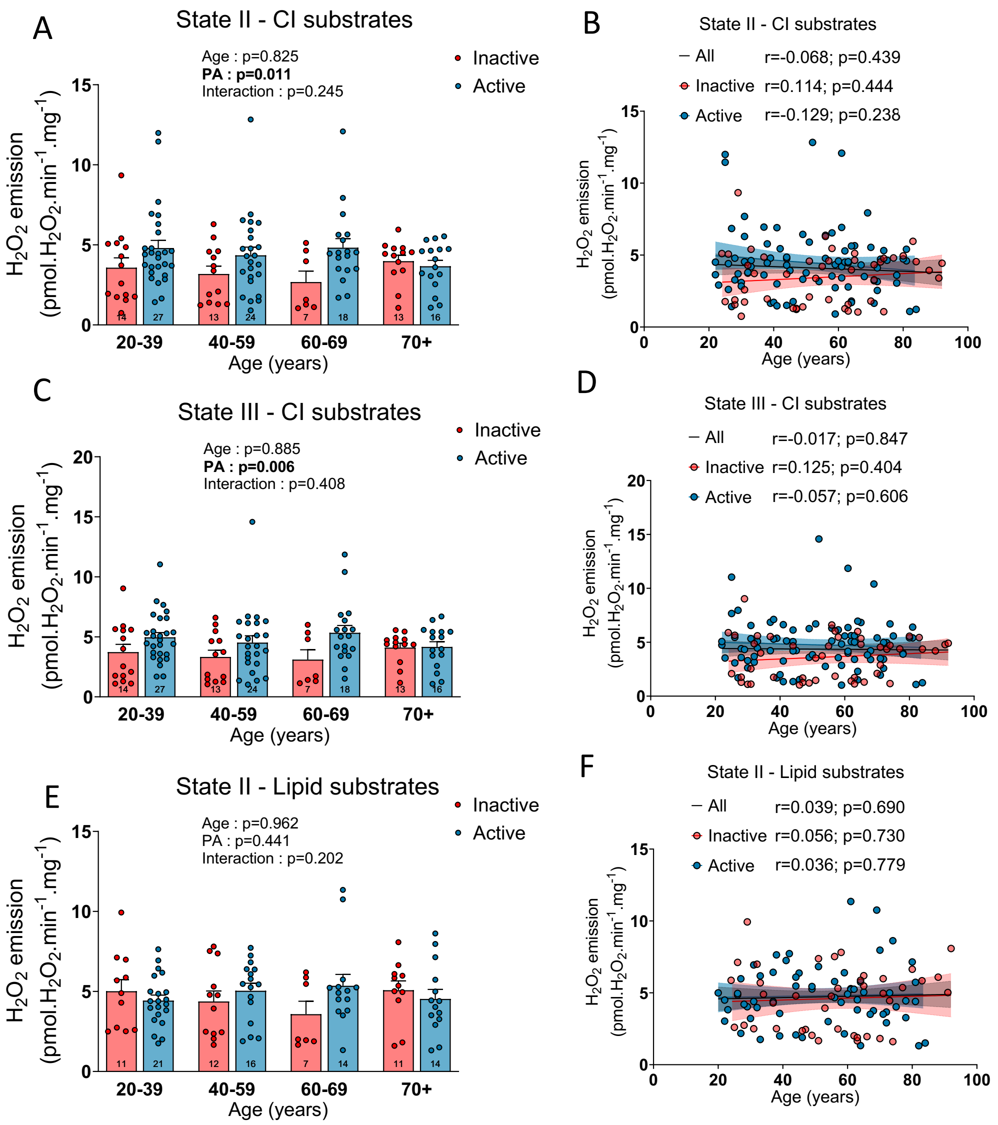


**Figure S6: Supplementary data on the impacts of aging and physical activity status on mitochondrial $H_2O_2$ emission.**

(A-B) State II and (C-D) state III (ADP stimulated) $H_2O_2$ emission rates driven by complex I substrates (Glutamate + Malate) in inactive and active participants. (E-F) State II $H_2O_2$ emission rates driven by lipids substrates (Palmitoyl-L-carnitine and Malate). For group analyses, results of the two-way ANOVA are displayed above each bar graph. Tukey post hoc tests were performed to test differences between age groups. Differences between inactive and active participants within each age group was assessed using multiple bilateral t-tests with FDR correction. Groups that do not share the same letter are significantly different. Linear regressions were performed to assess associations between age and variables of interest in the entire cohort (all) and for inactive and active participants separately. Pearson correlation coefficient (r) and p-values are displayed above each scatter plot. For two-way ANOVA, post hoc testing, and regression analyses, $p<0.05$ was considered statistically significant. For FDR analyses, $q<0.05$ was considered statistically significant.

**Figure S7:** The impacts of aging and physical activity status on mitochondrial free radical leak.

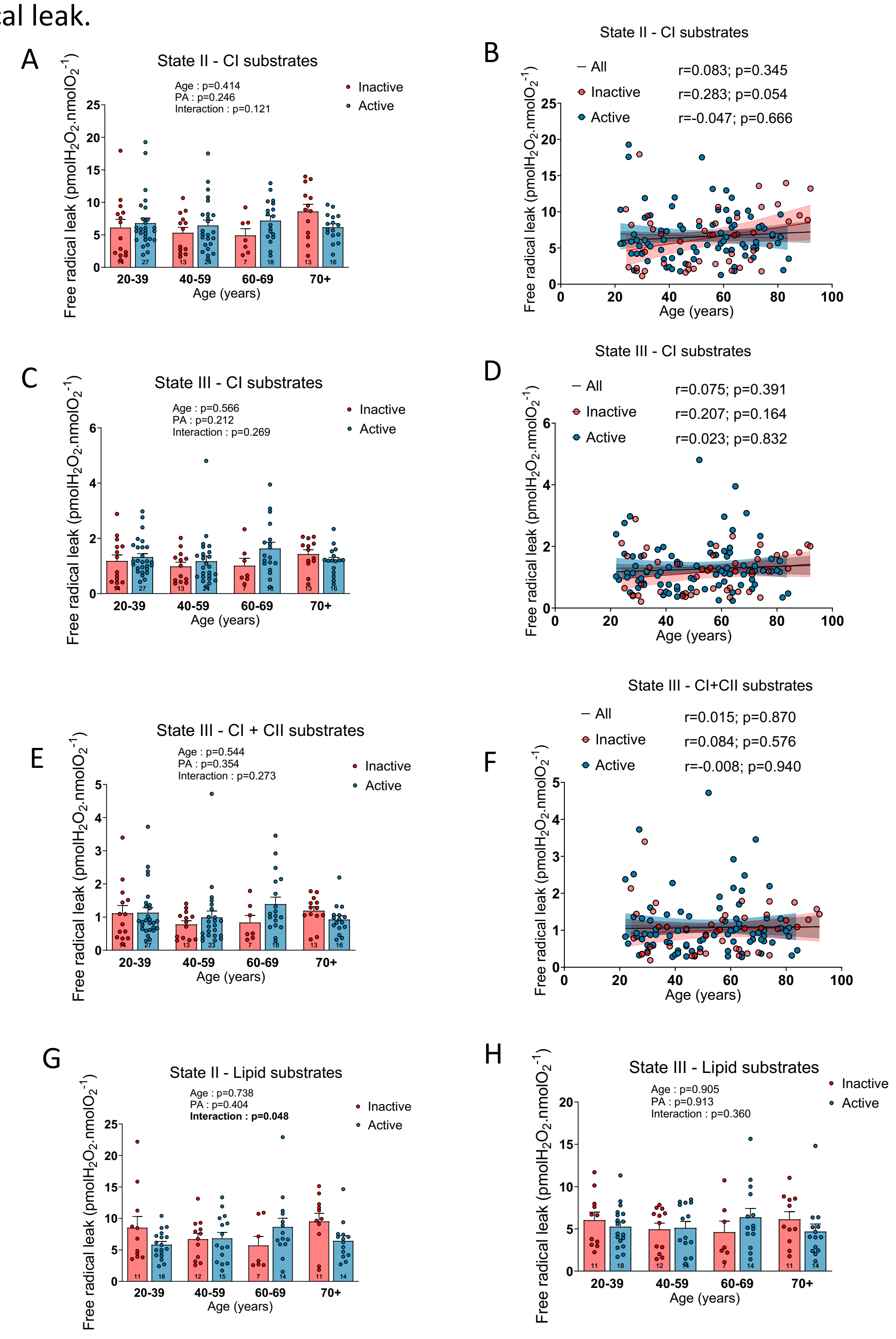


**Figure S7: The impacts of aging and physical activity status on mitochondrial free radical leak.**

(A-B) State II and (C-D) state III (ADP stimulated) free radical leak driven by complex I substrates (Glutamate + Malate) in inactive and active participants. (E-F) State III (ADP stimulated) free radical leak driven by complex I (Glutamate + Malate) and complex II

(Succinate) substrates. (G) State II and (H) State III (ADP stimulated) free radical leak driven by lipids substrates (Palmitoyl-L-carnitine and Malate). For group analyses, results of the two-way ANOVA are displayed above each bar graph. Tukey post hoc tests were performed to test differences between age groups. Differences between inactive and active participants within each age group was assessed using multiple bilateral t-tests with FDR correction. Groups that do not share the same letter are significantly different. Linear regressions were performed to assess associations between age and variables of interest in the entire cohort (all) and for inactive and active participants separately. Pearson correlation coefficient (r) and p-values are displayed above each scatter plot. For two-way ANOVA, post hoc testing, and regression analyses, $p<0.05$ was considered statistically significant. For FDR analyses, $q<0.05$ was considered statistically significant.

**Figure S8**: Correlations between state III $H_2O_2$ emission, physical function and muscle area

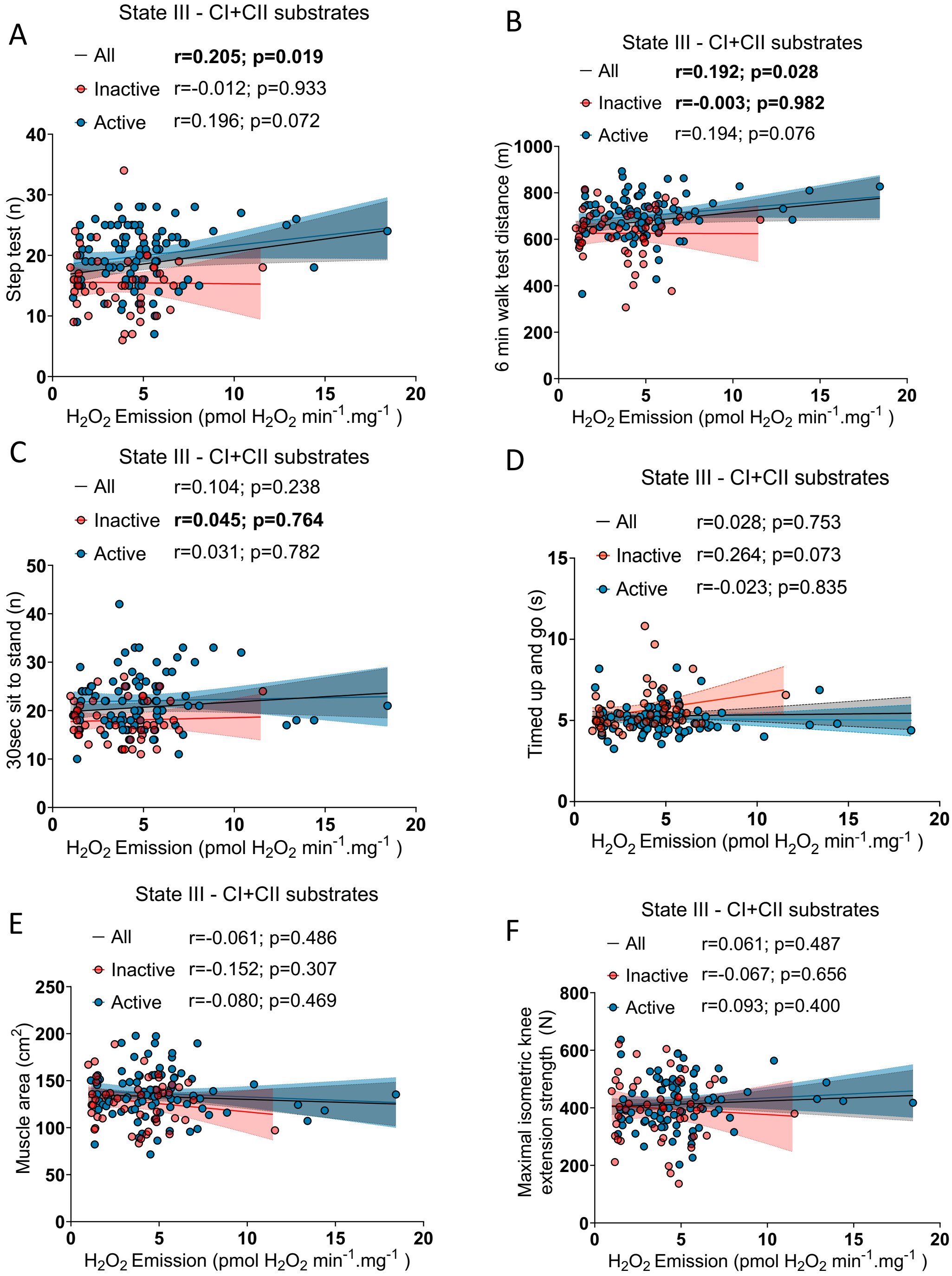


**Figure S8: Correlations between state III $H_2O_2$ emission, physical function and muscle area**

Correlations between state III (ADP stimulated) $H_2O_2$ emission rates driven by complex I (Glutamate + Malate) and II (Succinate) substrates and (A) step test, (B) 6 min walk test distance, (C) sit to stand test (number of repetitions performed in 30s), (D) timed up and go, (E) muscle cross-sectional area and (F) maximal isometric knee extension strength in inactive and active participants. Linear regressions were performed to assess associations between age and variables of interest in the entire cohort (all) and for inactive and active participants separately. Pearson correlation coefficient (r) and p-values are displayed above each scatter plot. p<0.05 was considered statistically significant.

**Figure S9:** Correlations between maximal $H_2O_2$ emission, physical function and muscle area

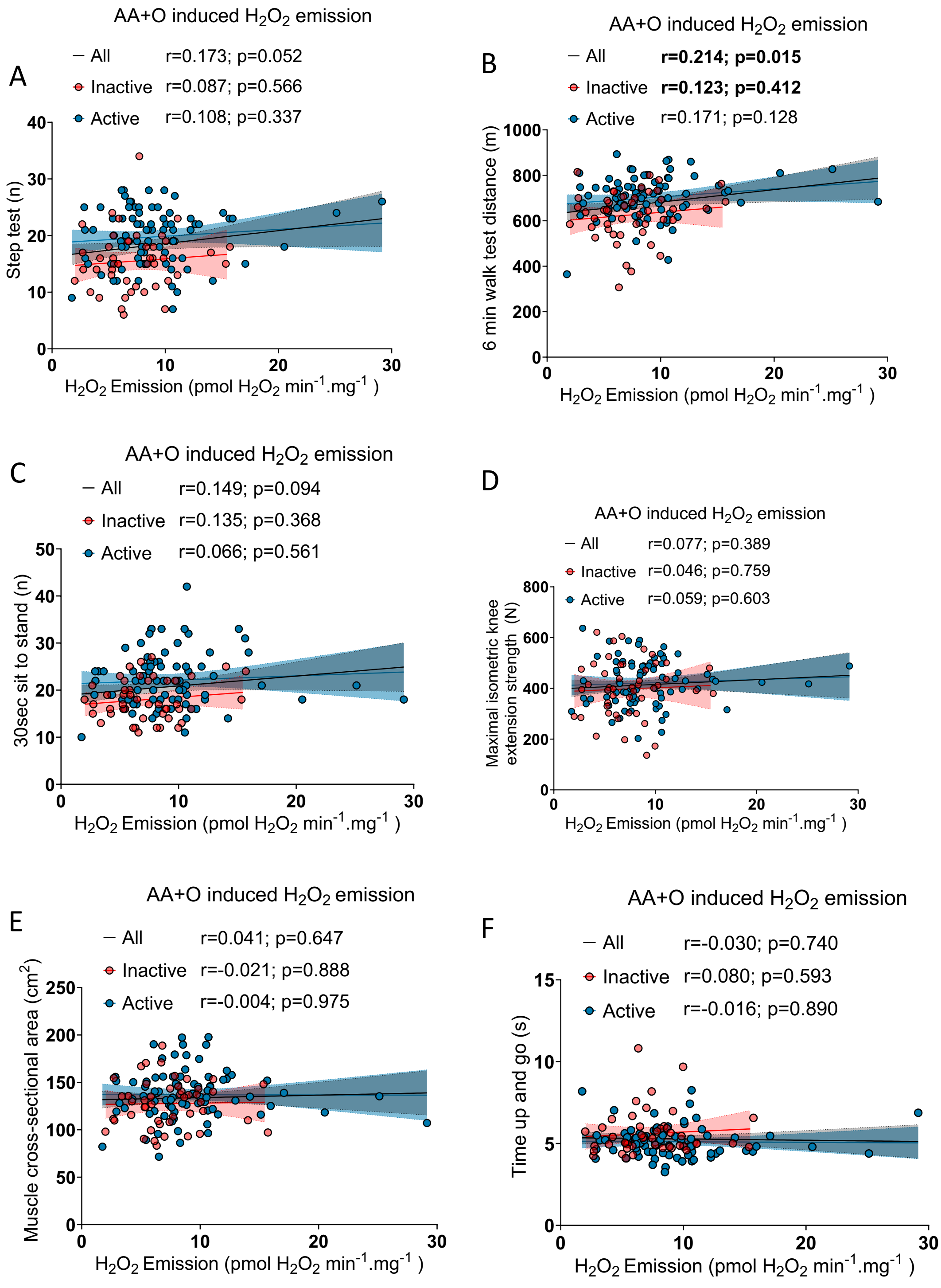

**Figure S9: Correlations between maximal $H_2O_2$ emission, physical function and muscle area**

Correlations between maximal $H_2O_2$ emission rates induced by antimycin A (AA) and oligomycin (O) and (A) step test, (B) 6 min walk test distance, (C) sit to stand test (number of repetitions performed in 30s), (D) timed up and go, (E) muscle cross-sectional area and (F) maximal isometric knee extension strength in inactive and active participants. Linear regressions were performed to assess associations between age and variables of interest in the entire cohort (all) and for inactive and active participants, separately. Pearson correlation coefficient (r) and p-values are displayed above each scatter plot. $p<0.05$ was considered statistically significant.

**Figure S10:** Relationships between mitochondrial calcium retention capacity, time to mPTP opening, muscle cross-sectional area and age.

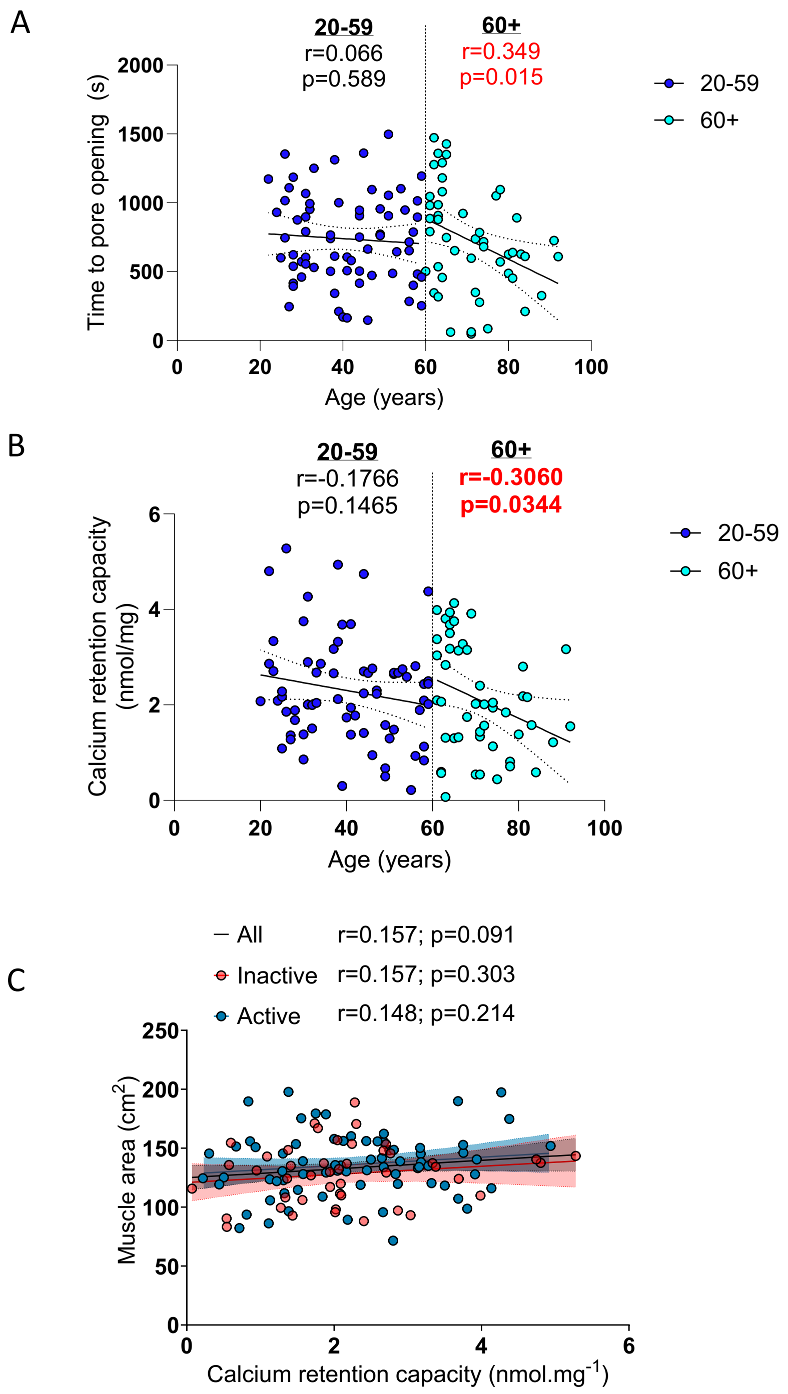


**Figure S10: The impacts of aging and physical activity status on mitochondrial calcium retention capacity and time to permeability transition pore opening**

Correlations between (A) time to permeability transition pore opening, (B) mitochondrial calcium retention capacity and age for all participants. Correlations between (C) mitochondrial calcium retention capacity and muscle cross-sectional area in inactive and active participants.

Linear regressions were performed to assess associations between age and variables of interest in the entire cohort (all) and for inactive and active participants separately. Pearson correlation coefficient (r) and p-values are displayed above each scatter plot. For two-way ANOVA, post hoc testing, and regression analyses, p<0.05 was considered statistically significant. For FDR analyses, q<0.05 was considered statistically significant. *=q<0.05.

**Figure S11:** The impacts of aging and physical activity status on mitochondrial calcium uptake rate.

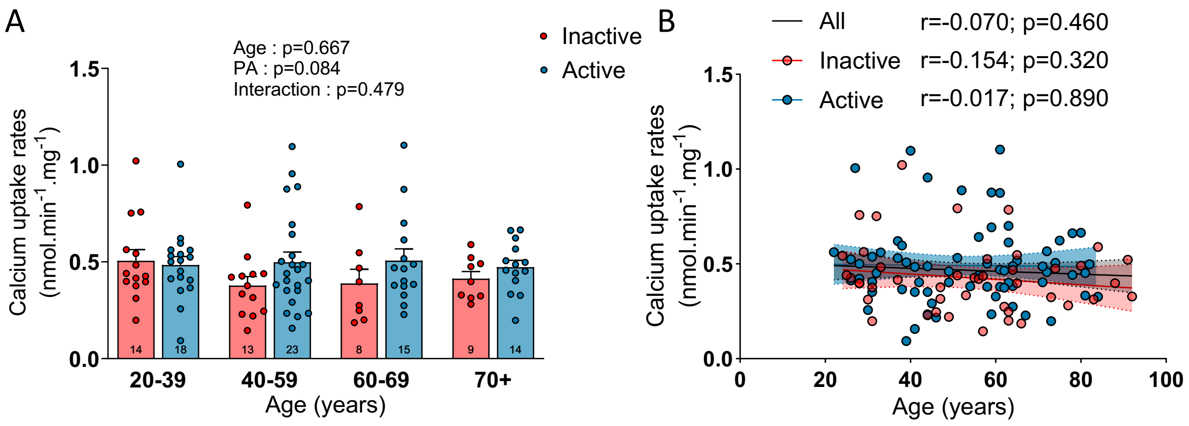


**Figure S11: The impacts of aging and physical activity status on mitochondrial calcium uptake rate.**

(A-B) Calcium uptake rates in inactive and active participants. For group analyses, results of the two-way ANOVA are displayed above each bar graph. Tukey post hoc tests were performed to test differences between age groups. Differences between inactive and active participants within each age group was assessed using multiple bilateral t-tests with FDR correction. Groups that do not share the same letter are significantly different. Linear regressions were performed to assess associations between age and variables of interest in the entire cohort (all) and for inactive and active participants separately. Pearson correlation coefficient (r) and p-values are displayed above each scatter plot. For two-way ANOVA, post hoc testing, and regression analyses, p<0.05 was considered statistically significant. For FDR analyses, q<0.05 was considered statistically significant. *=q<0.05.

**Figure S12:** Correlations between plasma GDF15 levels, age, mitochondrial respiration, and mitochondrial $H_2O_2$ emission.

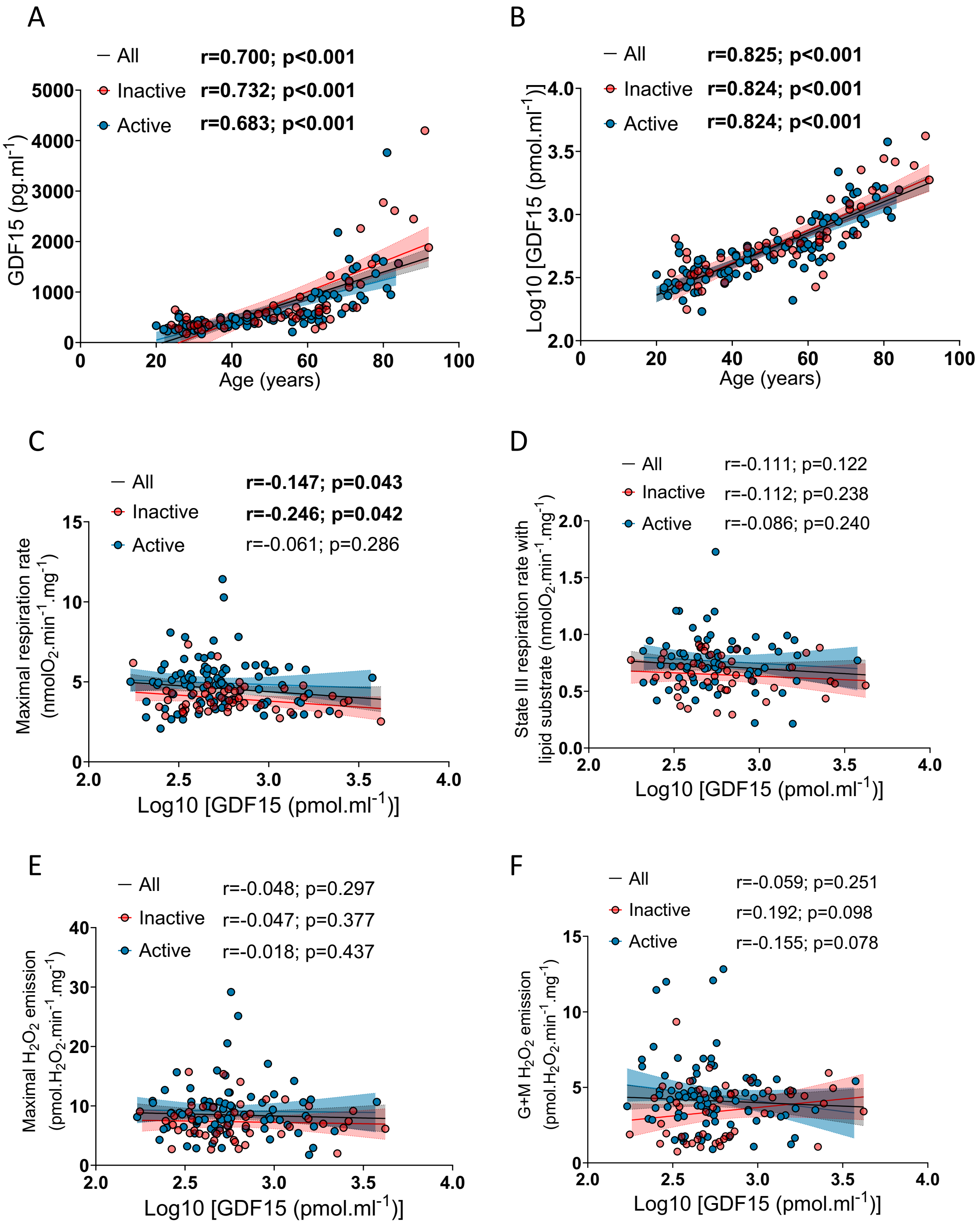


**Figure S12: Correlations between plasma GDF15 levels, age, mitochondrial respiration, and mitochondrial $H_2O_2$ emission.**

Correlations between plasma GDF15 levels and age (A: raw GDF15 values; B: log transformed GDF15 values) state III (ADP stimulated) respiration rate driven by complex I (Glutamate +

Malate) and II (Succinate) substrates (C, GDF15 values displayed on a log10 scale), state III (ADP stimulated) respiration rate driven by lipid substrates (palmitoyl-l-carnitine + malate) (D, GDF15 values displayed on a log10 scale), maximal $H_2O_2$ emission rates induced by antimycin A and oligomycin (E, GDF15 values displayed on a log10 scale), and state II $H_2O_2$ emission rates driven by complex I substrates (Glutamate + Malate) (E, GDF15 values displayed on a log10 scale), in inactive and active participants. Linear regressions were performed to assess associations between age and variables of interest in the entire cohort (all) and for inactive and active participants separately. Pearson correlation coefficient (r) and p-values are displayed above each scatter plot. $p<0.05$ was considered statistically significant.
